# A *cis*-regulatory element regulates *ERAP2* expression through autoimmune disease risk SNPs

**DOI:** 10.1101/2023.03.03.530973

**Authors:** Wouter J. Venema, Sanne Hiddingh, Jorg van Loosdregt, John Bowes, Brunilda Balliu, Joke H. de Boer, Jeanette Ossewaarde-van Norel, Susan. D. Thompson, Carl D. Langefeld, Lars T. van der Veken, Konstantinos Sofiadis, Peter H.L. Krijger, Wouter de Laat, Jonas J.W. Kuiper

## Abstract

Single nucleotide polymorphisms (SNP) near the *ERAP2* gene are associated with autoimmune conditions such as *Crohn’s disease*, and *birdshot chorioretinopathy*, as well as protection against lethal infections, including the *Black Death*. Due to high linkage disequilibrium (LD), a great number of trait-associated SNPs are correlated with *ERAP2* expression, however their functional mechanisms remain unidentified. We used genome editing and functional genomics to identify causal variants that remain obscured by LD. We demonstrate by reciprocal allelic replacement that *ERAP2* expression is directly controlled by the genotype of splice region SNP rs2248374. However, we demonstrate that autoimmune disease-risk SNPs located near the downstream *LNPEP* gene promoter are independently associated with *ERAP2* expression. Allele-specific conformation capture assays revealed long-range chromatin contacts between the *LNPEP* promoter region and the *ERAP2* promoter and showed that interactions were stronger in patients carrying the alleles that increase susceptibility to autoimmune diseases. Replacing the disease-associated SNPs in the *LNPEP* promoter by reference sequences lowered *ERAP2* expression. These findings show that clustered GWAS signals associated with diverse autoimmune conditions and lethal infections act in concert to control ERAP2 expression and that disease-associated variants can convert a gene promoter region into a potent enhancer of a distal gene.

## Background

MHC class I molecules (MHC-I) display peptides derived from intracellular proteins allowing CD8+ T cells to detect infection and malignancy (1, 2). In the endoplasmic reticulum, aminopeptidases ERAP1 and ERAP2 shorten peptides that are presented by MHC-I (3–5). Dysfunctional ERAP may alter the repertoires of peptides presented by MHC-I, potentially activating CD8+ T cells and causing adverse immune responses (6–8).

In *genome-wide association studies* (GWAS), polymorphisms at *5q15* (chromosome 5, *q* arm, G-band *15*) near the *ERAP1* and *ERAP2* genes have been associated with multiple autoimmune conditions. Among them are ankylosing spondylitis (9, 10), Crohn’s disease [CD] (11), juvenile idiopathic arthritis [JIA] (12), birdshot chorioretinopathy [BCR] (13, 14), psoriasis, and Behcet’s disease (15, 16). The SNPs identified in GWAS as disease-risk SNPs in *ERAP1* usually correspond to changes in amino acid residues, resulting in proteins with different peptide trimming activities and expression levels (8,17–20).

On the other hand, many SNPs near *ERAP2* are highly correlated with the level of *ERAP2* expression (i.e., *expression quantitative trait loci* [eQTLs] for *ERAP2*) (21, 22). Due to linkage disequilibrium (LD) between these SNPs, there are two common *ERAP2* haplotypes; one haplotype encodes enzymatically active ERAP2 protein while the alternative haplotype encodes transcript with an extended exon 10 that contains premature termination codons, inhibiting mRNA and protein expression (23). The haplotype that produces full-size *ERAP2* increases the risk of autoimmune diseases such as CD, JIA, and BCR, but it also protects against severe respiratory infections like pneumonia (24), as well as historically the *Black Death*, caused by the bacterium *Yersinia pestis* (11-13,25). There is a SNP rs2248374 (allele frequency ∼50%) located within a donor splicing site directly after exon 10 that tags these common haplotypes (14,23,26). Consequently, rs2248374 is assumed to be the sole variant responsible for ERAP2 expression. Although this is supported by association studies and minigene based assays (23, 26), strikingly, there have been no studies evaluating *ERAP2* expression after changing the allele of this SNP in genomic DNA. This leaves the question of whether the rs2248374 genotype is essential for *ERAP2* expression unanswered.

More than a hundred additional *ERAP2* eQTLs located in and downstream of the *ERAP2* gene, form a large ‘extended ERAP2 haplotype’ (13). It is commonly assumed that these *ERAP2* eQTLs work solely by tagging (i.e., in LD with) rs2248374 (23,25,27–30). There is however, evidence that some SNPs in the extended *ERAP2* haplotype may influence *ERAP2* expression independent of rs2248374 (20,32,32). The use of CRISPR-Cas9 genome editing and functional genomics may be able to unravel the *ERAP2* haplotypes and identify causal variants that regulate *ERAP2* expression but are obscured by LD with rs2248374 in association studies.

We investigated whether rs2248374 is sufficient for the expression of *ERAP2*. Polymorphisms influencing *ERAP2* expression were identified using allelic replacement by CRISPR-mediated homologous repair and conformation capture assays. We report that rs2248374 was indeed critical for *ERAP2* expression, but that *ERAP2* expression is further influenced by additional SNPs that facilitate a local conformation that increases promoter interactions.

## Results

### ERAP2 expression depends on the genotype of rs2248374

The SNP rs2248374 is located downstream of exon 10 of *ERAP2* and its genotype strongly correlates with ERAP2 expression. Predictions by deep neural network-based algorithms *SpliceAI* and *Pangolin* indicate that the A>G allelic substitution by rs2248374 inhibits constitutive splicing three base pairs upstream at the canonical exon-intron junction (*SpliceAI,* donor loss Δ score = 0.51, *Pangolin* Δ score = 0.58). Despite widespread assumption that this SNP controls *ERAP2* expression, functional studies are lacking (23). Therefore, we first aimed to determine whether ERAP2 expression is critically dependent on the genotype of this SNP. Allelic replacement by CRISPR-mediated homologous repair using a donor DNA template was used to specifically mutate rs2248374 G>A by homology directed repair (HDR) (**Figure 1A**). Because HDR is inefficient (42), a silent mutation was inserted into the donor template to produce a *Taql* restriction site, which can be used to screen clones with correctly edited SNPs. As THP-1 cells are homozygous for the G allele of rs2248374 (**Figure 1B**), we used this cell line for experiments because it can be grown in single-cell derived clones. We targeted rs2248374 in THP-1 cells and established a clone that was homozygous for the A allele of rs2248374 (**Figure 1B**). Sequencing of the junctions confirmed that the integrations were seamless and precisely positioned in-frame.

**Figure 1:**
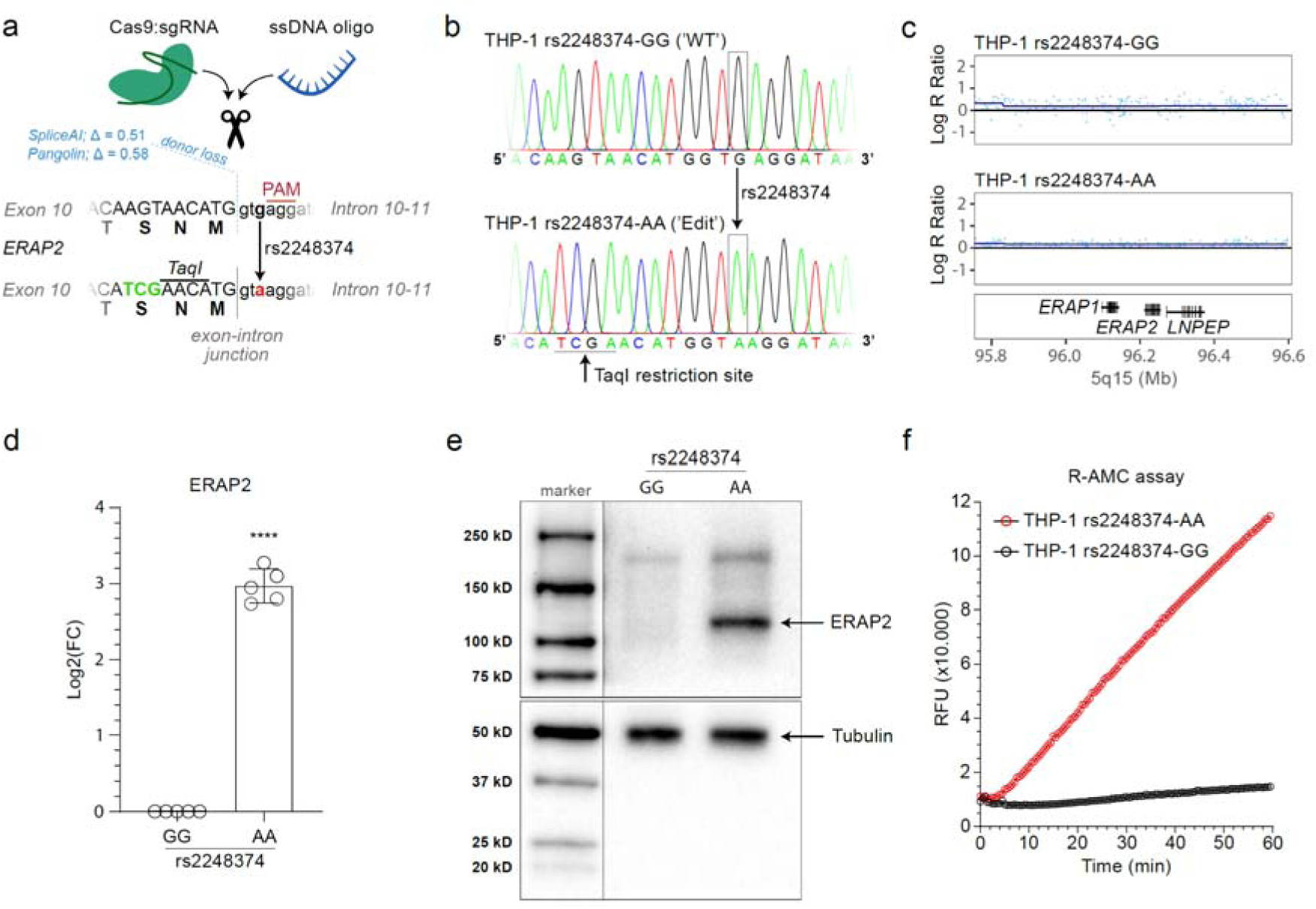
The A allele of rs2248374 is essential for full-length ERAP2 expression. **A**) Overview of the CRISPR-Cas9 mediated homology-directed-repair (HDR) strategy for SNP allelic replacement of the G allele of rs2248374 to the A allele in THP-1 cells. The ssDNA oligo template introduces the A allele at position rs2248374 and a silent *TaqI* restriction site used for screening successfully edited clones. The predicted effect size (delta scores from *SpliceAI* and *Pangolin*) and intended position that exhibits altered splicing induced by the G allele of rs2248374 is shown in blue. **B**) Sanger sequence data showing THP-1 ‘WT’ with the single rs2248374-G variant and the successful SNP modification to the A allele of rs2248374. **C**) SNP-array based copy number profiling and analysis of regions of homozygosity of unedited and edited THP-1 clones demonstrating no other genomic changes. Plot is zoomed on *5q15.* Genome-wide results are outlined in Supplemental Figure 3. **D**) *ERAP2* gene expression determined by qPCR in cellular RNA from THP-1 cells unedited or edited for the genotype of rs2248374. **E**) Western blot analysis of ERAP2 protein in cell lysates from THP-1 cells unedited or edited for the genotype of rs2248374. ****) t.test, *P*<0.001 **F**) Hydrolysis [expressed as relative fluorescence units (RFU)] of the substrate L-Arginine-7-amido-4-methylcoumarin hydrochloride (R-AMC) by immunoprecipitated ERAP2 protein from THP-1 cell lines unedited or edited for the genotype of rs2248374. The generation of fluorescent AMC indicates ERAP2 enzymatic activity.

SNP-array analysis was performed to exclude off-target genomic alterations giving rise to duplications and deletions in the genome of the gene edited cell lines (**Supplemental Figure 3**). We did not observe any of such unfavourable events. This confirmed that our editing strategy did not induce widespread genomic changes (43). While THP-1 cells are characterised by genomic alterations, including large regions of copy number neutral loss of heterozygosity of chromosome 5 (including *5q15*) (44, 45) (**Supplemental Figure 3**), the results confirmed that single-cell clones from the unedited ‘wild-type’ (WT, rs2248374-GG) THP-1 cells and "edited" THP-1 (rs2248374-AA) were genetically identical at *5q15*, which justifies their comparison (**Figure 1C**). In contrast with WT THP-1, *ERAP2* transcript became well detectable in THP-1 cells in which we introduced the A allele of rs2248374 (**Figure 1D**). According to Western Blot analysis, WT THP-1 cells lack ERAP2 protein, while the rs2248374-AA clone expressed full-length ERAP2 (**Figure 1E**), which was enzymatically functional as determined by a fluorogenic *in vitro* activity assay (**Figure 1F**).

Oppositely, we then examined whether mutation of rs2248374 A>G would abolish *ERAP2* expression in cells naturally expressing ERAP2. The Jurkat T cell line was chosen because they are heterozygous for rs2248374 and express ERAP2, and they possess the ability to grow in single-cell clones required to overcome the low-efficiency of CRISPR knock-in by HDR. To alter the single A allele of rs2248374 in the Jurkat cell line, we used a donor DNA template encoding the G variant (**Supplemental Figure 4A**) and established a clone homozygous for rs2248374 G (**Supplemental Figure 4B**). We found no changes between our unedited population and rs2248374 edited Jurkat cells at *5q15* by whole genome homozygosity mapping (**Supplemental Figure 4C**). The A>G substitution at position rs2248374 depressed *ERAP2* mRNA expression (**Supplemental Figure 4D**) and abolished ERAP2 protein expression (**Supplemental Figure 4E**). These results show that *ERAP2* mRNA and protein expression is critically dependent on the genotype of rs2248374 at steady-state conditions.

### Disease risk SNPs are associated with ERAP2 levels independent of rs2248374

Many additional SNPs at chromosome *5q15* show strong associations with *ERAP2* gene expression levels (also known as *ERAP2* eQTLs). Despite LD between the other *ERAP2* eQTLs and rs2248374, rs2248374 does not appear to be the strongest *ERAP2* eQTL in the GTEx database (data for GTEx ‘whole blood’ shown **Figure 2A, Supplementary Table 2**). Following this, we investigated the SNPs near the *ERAP2* gene that are associated with several T-cell-mediated autoimmune conditions, such as CD, JIA, and BCR (**Figure 2B-D**). We found strong evidence for colocalization between GWAS signals at *5q15* for BCR, CD, and JIA and *cis*-eQTLs for *ERAP2* (**Supplementary Table 3-5**) (posterior probability of colocalization >90%). This indicates that these SNPs alter the risk for autoimmunity through their effects on *ERAP2* gene expression. It is noteworthy, however, that the GWAS hits at *5q15* for CD, BCR, and JIA are in high LD (r^2^>0.9) with each other but not in high LD with rs2248374 (r^2^<0.8) (**Figure 2E**). Furthermore, the GWAS association signal at *5q15* for JIA (lead variant rs27290; *P*_dominant_= 7.5 × 10^-9^) did not include rs2248374 (JIA, *P*_dominant_ = 0.65) (**Figure 2D**). In line with this, we previously reported that the lead variant rs7705093 (**Figure 2C**) is associated with BCR after conditioning on rs2248374 (31). These findings reveal that SNPs implicated in these complex human diseases by GWAS may affect *ERAP2* expression through mechanisms other than rs2248374.

**Figure 2:**
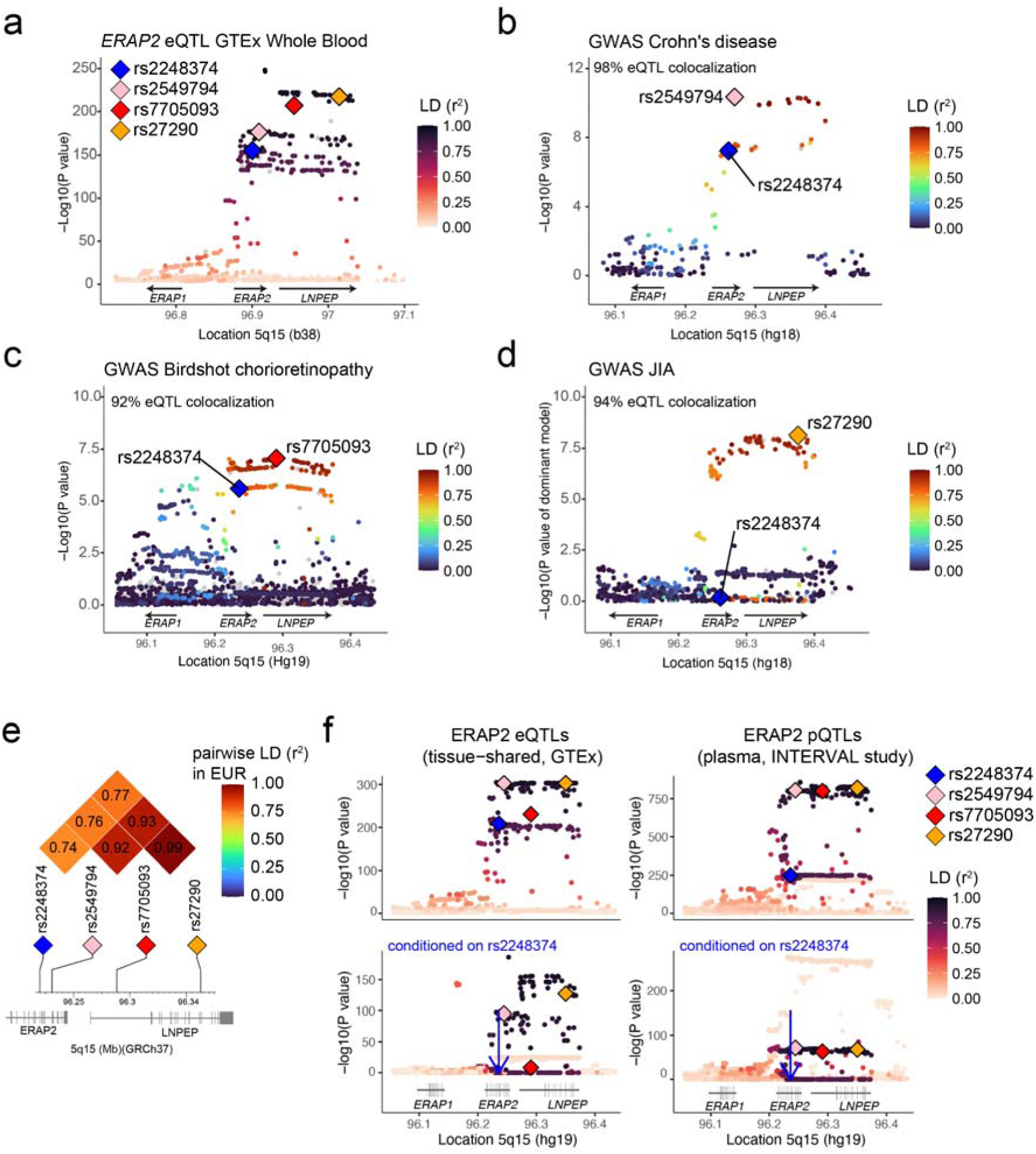
Autoimmune disease-risk SNPs associated with ERAP2 levels independent from rs2248374 genotype. **A**) *ERAP2* eQTL data from GTEx whole blood (**Supplementary Table 2**). GWAS lead variants at *5q15* for Crohn’s disease [CD] (rs2549794, see *b*), birdshot chorioretinopathy [BCR] (rs7705093, see *c*), and juvenile idiopathic arthritis [JIA] (rs27290, see *d*) and rs2248374 are denoted by coloured diamonds. The colour intensity of each symbol reflects the extent of LD (r^2^ from 1000 Genomes EUR samples) with rs2927608. Grey dots indicate missing LD information. **B-D**) Regional association plots of GWAS from CD, BCR, and JIA (**Supplementary Table 3-5**). For the CD we used the *P* value of rs2549782 (LD [r2] =1.0 with rs2248374 in EUR). The colour intensity of each symbol reflects the extent of LD (r^2^ estimated using 1000 Genomes EUR samples) with rs2927608. The results from colocalization analysis between GWAS signals and *ERAP2* eQTL data from whole blood (in *a*) is denoted (details in Appendix I). **E**) Pairwise LD (r^2^ estimated using 1000 Genomes EUR samples) comparison between splice variant rs2248374 (*ERAP2*), and disease-risk SNPs rs2549794 (CD), rs7705093 (BCR) and rs27290 (JIA). **F**) Initial association results and conditional testing of *ERAP2* tissue-shared eQTL data from GTEx consortium and ERAP2 pQTL data from plasma proteomics of the INTERVAL study (**Supplementary Table 6 and 7**) (37). Conditioning on rs2248374 (dark blue diamond) revealed independent *ERAP2* eQTL and ERAP2 pQTL signals that include lead variants at *5q15* for CD, BCR, and JIA (*P* <5.0 x 10^-8^). The human reference sequence genome assembly annotations are indicated.

We therefore sought to determine if *ERAP2* eQTLs function independently of rs2248374. In agreement with the role ERAP2 plays in the MHC-I pathway that operates in most cell types, *ERAP2* eQTLs are shared across many tissues (46, 47). As a proof of principle, we used tissue-shared *ERAP2* eQTLs from a previous analysis of RNA-seq data from the GTEx Consortium (**Figure 2F**). To test whether the disease-associated top association signals were independent from rs2248374, we performed conditional testing of the *ERAP2* eQTL signal by first including the genotype of rs2248374 as a covariate in the regression model. Conditioning on rs2248374 revealed a complex independent *ERAP2* eQTL signal composed of many SNPs extending far downstream into the *LNPEP* gene (**Figure 2F**). This secondary *ERAP2* eQTL signal included the lead variants at *5q15* for CD, BCR, and JIA (*P_conditioned_* < 5.9 × 10^-9^), consistent with earlier findings in association studies (**Supplemental Table 6**) (20, 31). We further strengthened these observations by using summary statistics from SNPs associated with plasma levels of ERAP2 from the INTERVAL study (called protein quantitative trait loci, or pQTLs)(37). In agreement with the mRNA data from GTEx, conditioning on rs2248374 revealed a strong independent association between GWAS lead variants and ERAP2 protein levels (*P_conditioned_* < 1.6 × 10^-62^) (**Figure 2F**). Based on these results, we conclude that GWAS signals at *5q15* are associated with ERAP2 levels independently of rs2248374.

### SNPs in a downstream cis-regulatory element modulate ERAP2 promoter interaction

Computational tools to predict the functional impact of non-coding variants may be highly inaccurate (48). To prioritise likely causal variants by experimentally monitoring their effects on ERAP2, we aimed to resolve the function of SNPs that correlated with ERAP2 expression independent from rs2248374. First, we aimed to generate a large 116 Kb heterozygous deletion downstream of *ERAP2* in *Jurkat* cells. This deletion included all downstream SNPs as well as the entire *LNPEP* gene (**Supplemental Figure 1**). *ERAP2* expression by qPCR was not significantly reduced by this approach, which may be due to only partial deletion of the region as shown by whole genome zygosity mapping (**Supplemental Figure 1**), or the non-selective removal of all downstream putative regulatory elements.

Since allelic replacement would provide a more physiological relevant approach, we next aimed to specifically alter the SNP alleles and evaluate the impact on *ERAP2* expression. The large size of the region containing all the “independent” *ERAP2* eQTLs prevents efficient HDR (43), so we decided to prioritise a regulatory interval with *ERAP2* eQTLs. Genetic variation in non-coding enhancer sequences near genes can influence gene expression by interacting with the gene promoter (49). Therefore, we leveraged chromosome conformation capture coupled with sequencing (Hi-C) data enriched by chromatin immunoprecipitation for the activating histone H3 lysine 27 acetylation (*H3K27ac*, an epigenetic mark of active chromatin that marks enhancer regions) in primary T cells, B cells, and monocytes (40), immune cells that share ERAP2 eQTLs as shown by single-cell sequencing studies (47). We selected *ERAP2* eQTLs located in active enhancer regions at *5q15* (i.e., H3K27ac peaks) that significantly interacted with the transcriptional start site of *ERAP2* for each immune cell type. This revealed diverse and cell-specific significant interactions of *ERAP2* eQTLs across the extended *ERAP2* haplotype in immune cells, indicating many regions harbouring eQTLs that were physically in close proximity with the transcription start site of *ERAP2* (**Figure 3A**). Note that none of these SNPs showed significant interaction with the promoters of *ERAP1* or *LNPEP*. Among these, 9 common non-coding SNPs concentrated in a ∼1.6 kb region downstream of *ERAP2* at the 5‘end of the gene body of *LNPEP* exhibited strong interactions with the *ERAP2* promoter (**Figure 3A**), suggesting that these SNPs lie within a potential regulatory element (i.e., enhancer) that is active in multiple cell lineages. Consistent with these data, examination of ENCODE data of heart, lung, liver, skeletal muscle, kidney, and spleen revealed enrichment of *H3K27ac* marks spanning the 1.6 kb locus, supporting that these SNPs lie within an enhancer-like DNA sequence that is active in across tissues (**Figure 3A**). This also corroborates the finding that these SNPs are *ERAP2* eQTLs across tissues, as we (36) showed previously (**Figure 2F**). Results from a recent targeted massively parallel reporter assay (MPRA) (32) support that this region may exhibit differential regulatory effects (i.e., altered transcriptional regulation), depending in particular on the allele of SNP rs2548224 (difference in expression levels of target region; reference versus alternative allele for rs2548224, *Padj* = 4.9 × 10^-3^)(**Figure 3B**). This SNP is also a very strong (rs2248374-independent) *ERAP2* eQTL and pQTL (**Supplemental Figure 5, Supplementary Table 1**). In light of the fact that this selected region downstream of *ERAP2* contained SNPs which are associated with ERAP2 expression independently of rs2248374, is physically in close proximity with the *ERAP2* promoter (i.e., by Hi-C), and may exert allelic-dependent effects (i.e., by MPRA), we hypothesised that the risk alleles of these SNPs associated with autoimmunity may increase the interaction with the promoters of *ERAP2*.

**Figure 3:**
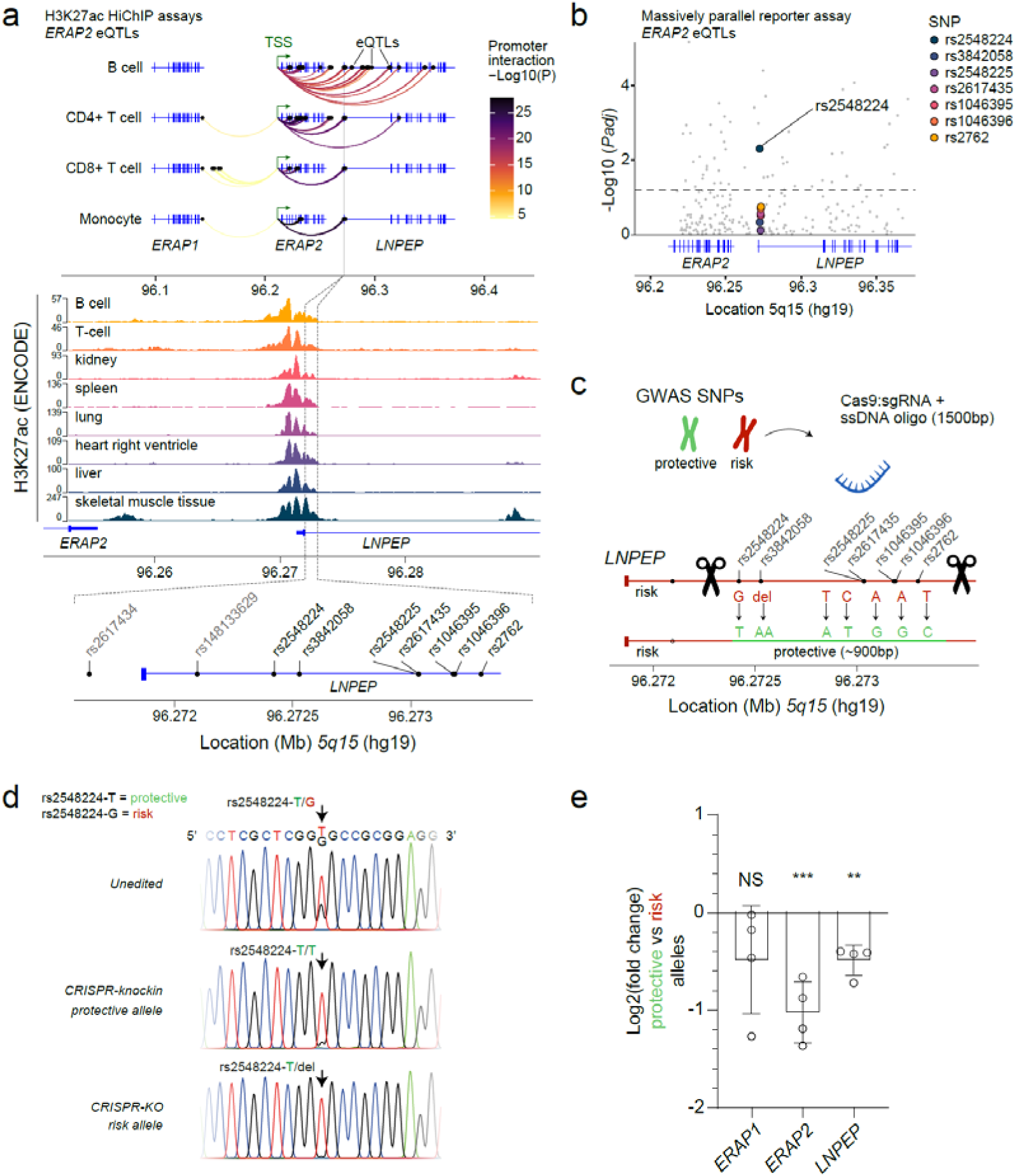
Autoimmune disease risk SNPs tag a downstream regulatory element that regulates *ERAP2* expression. **A**) Chromosome conformation capture coupled with sequencing (Hi-C) data enriched by chromatin immunoprecipitation for the histone H3 lysine 27 acetylation (*H3K27ac*) in primary immune cells from Chandra *et al.* (40). Highlighted are the *ERAP2* eQTLs (black dots) that overlap with *H3K27ac* signals that significantly interact with the transcriptional start site of *ERAP2* in four different immune cell types (B cells, CD4+ T cells, CD8+ T cells and monocytes). Nine common non-coding SNPs concentrated in a ∼1.6kb region exhibited strong interactions and overlay with H3K27ac signals from ENCODE data of heart, lung, liver, skeletal muscle, kidney and spleen revealed. **B**) The -Log10 *P* values (adjusted for multiple testing using the Benjamini-Hochberg) of the effect of 986 *ERAP2* eQTLs on differential expression (alternative versus reference allele) of their 150bp window region from a massively parallel reporter assay as reported by Abell *et al.* (32). The 7 SNPs identified by HiChIP in *a* are colour-coded **C**) Overview of the homology-directed-repair (HDR) strategy to use CRISPR-Cas9 mediated SNP replacement in Jurkat cells to switch the alleles from disease risk SNPs (i.e., alleles associated with higher ERAP2 levels) to protective haplotype (i.e., alleles associated with lower ERAP2 expression). The region from 5’ to 3’ spans 879 bp. **D**) Sanger sequencing results for the genotype of rs2548224 for Jurkat cells targeted by the CRISPR-knockin approach outlined in *c.* in comparison to unedited Jurkat cells and Jurkat cells in which the risk haplotype was deleted by CRISPR-Cas9-mediated knock-out as shown in **Supplementary Figure 1**. **E**) Expression of *ERAP2*, *LNPEP* and *ERAP1* by qPCR in Jurkat clones after allelic substitution of rs2548224. Data represents n = 4 biological replicates, Two-tailed unpaired t.test was assessed to compare WT expression to the modified clone (** *P*<0.01, *** *P*<0.001).

To investigate this, we first asked if specific introduction of the alternative alleles for these SNPs would affect the transcription of *ERAP2*. We targeted this region of the *ERAP2*-encoding chromosome in Jurkat cells using CRISPR-Cas9 and two guide RNAs in the presence of a large (1500 bp) single-stranded DNA template identical to the target region, but encoding the alternative alleles for 7 of the 9 non-coding SNPs. These SNPs were selected, because these cluster close together (∼900 bp distance from 5’ SNP rs2548224 to 3’ SNP rs2762) and are in tight LD (r^2^∼1 in EUR) with each other, as well as with the GWAS lead variants at *5q15* from CD, BCR, and JIA (r^2^>0.9) (**Supplemental Figure 6**). The introduction of the template DNA for CRISPR-knock-in by HDR did not induce other genomic changes (**Figure 3C** and **Supplemental Figure 7**). Sanger sequencing revealed targeting this intronic region by CRISPR-mediated HDR successfully altered the allele for SNPs rs2548224 in the regulatory element, but not the other targeted SNPs (**Figure 3D**, and **Supplemental Figure 8**) Regardless, altering the risk allele T to the reference allele G for rs2548224 for this SNP resulted in significant decrease in *ERAP2* mRNA (unpaired t-test, *P* = 3.0 × 10^-4^) (**Figure 3E**). In agreement with the known ability of enhancers to regulate multiple genes within the same topologically associated domain, altering the alleles of these SNPs also resulted in significant reductions in the expression of the *LNPEP* gene, (unpaired t-test, *P* = 0.0018), but not *ERAP1* (**Figure 3E**). Overall, these results indicate that *ERAP2* gene expression is affected by disease-associated SNPs downstream of the *ERAP2* gene.

### ERAP2 promoter contact is increased by autoimmune disease risk SNPs

RegulomeDB indicates that the SNP rs2548224 overlapped with 153 epigenetic mark peaks in various cell types (e.g., *POL2RA* in B cells). Considering its position within *LNPEP*’s promoter region, it makes it difficult to distinguish between local promoter and enhancer functions. In agreement with the inefficiency of targeting intronic regions by CRISPR *knock-in*, efforts to introduce the risk alleles of SNP in THP-1 (rs2248374-AA clone in **Figure 1**) cells that carry the reference alleles for these SNPs were not successful. To determine whether alleles of the SNPs in the regulatory element directly influenced contact with the *ERAP2* promoter, we used allele-specific 4C-seq (50) in B cell lines generated from blood of three BCR patients carrying both the risk and non-risk allele (i.e., heterozygous for disease risk SNPs). Using nuclear proximity ligation, 4C-seq enables the quantification of contact frequencies between a genomic region of interest and the remainder of the genome (34). Allele-specific 4C-seq has the advantage of measuring chromatin contacts of both alleles simultaneously and allows comparison of the risk allele versus the protective allele in the same cell population. We found that the downstream regulatory region formed specific contacts with the promoter of *ERAP2* (**Figure 4A**). Moreover, in two out of three patients contact frequencies with the *ERAP2* promoter were substantially higher for the risk allele than the protective allele, supporting the idea that *ERAP2* upregulation may be a consequence of a direct regulatory interaction between the autoimmune risk SNPs and the gene promoter (**Figure 4B**).

**Figure 4:**
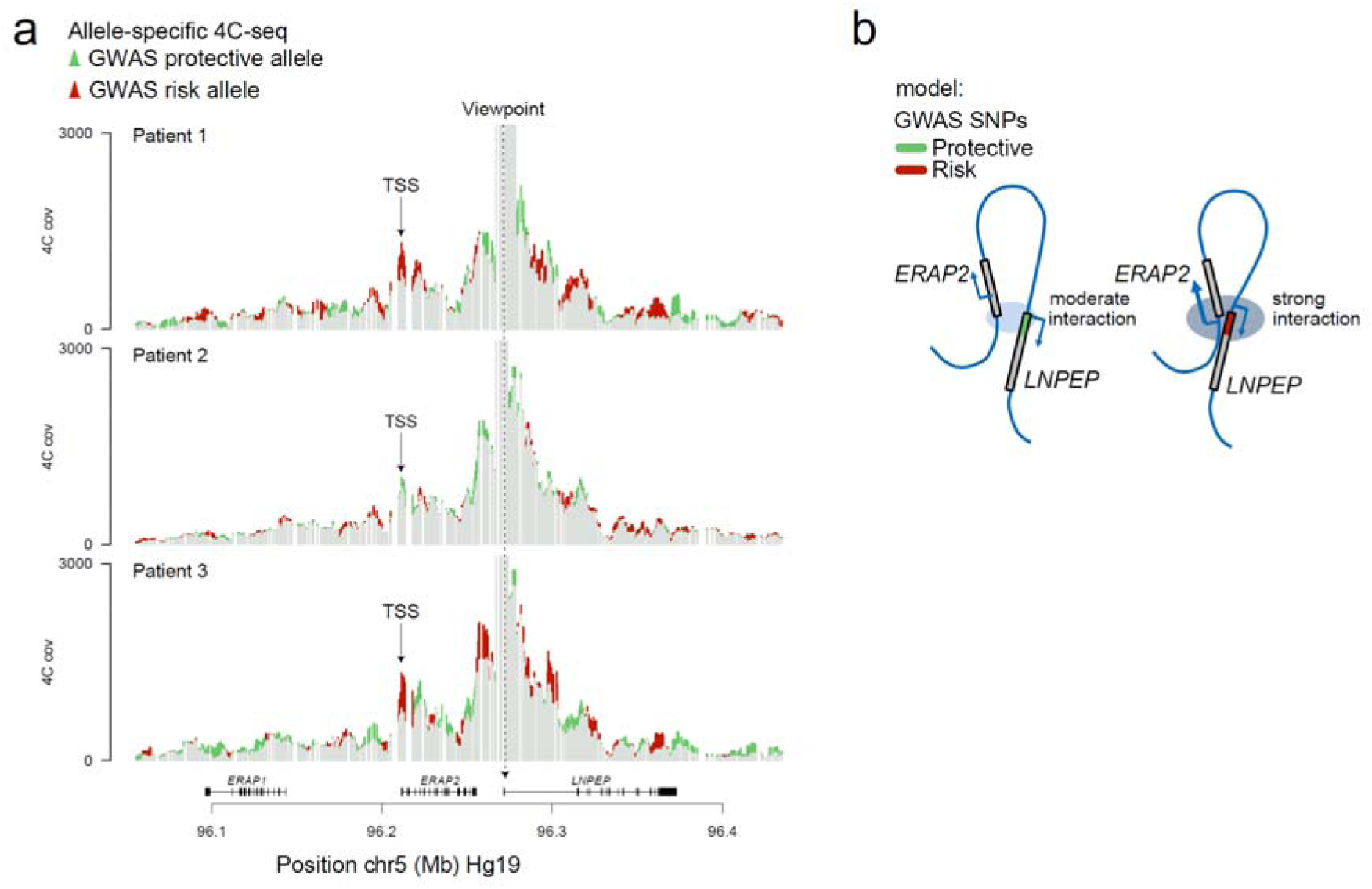
Autoimmune disease risk SNPs show high contact frequency with the *ERAP2* promoter in autoimmune patients. 4C analysis of contacts between the downstream regulatory region across the *ERAP2* locus. **A)** 4C-seq contact profiles across the *ERAP2* locus in B cell lines from three patients with BCR that are heterozygous (e.g., rs2548224-G/T) for the *ERAP2* eQTLs located in the downstream regulatory element (the 4C viewpoint is centred on the SNP rs3842058 in the *LNPEP* promoter as depicted by the dashed line). The Y-axis represents the normalised captured sequencing reads. The red lines in each track indicate the regions where the risk alleles show more interactions compared with the reference alleles while the green lines indicate the regions where the reference alleles (i.e., protective alleles) show more interactions. TSS = transcription start site of *ERAP2*. **B)** Schematic representation of ERAP2 regulation by autoimmune risk SNPs in the downstream regulatory element. showing the regulatory element with risk alleles (red), or reference (protective) alleles (green). The DNA region surrounding the *ERAP2* and *LNPEP* gene is shown in blue.

## Discussion

In this study, we demonstrated that ERAP2 expression is initiated or abolished by the genotype of the common SNP rs2248374. Furthermore, we demonstrated that autoimmune disease risk SNPs identified by GWAS at *5q15* are statistically associated with *ERAP2* mRNA and protein expression independently of rs2248374. We show that autoimmune risk SNPs tag a gene-proximal DNA sequence that influences *ERAP2* expression and interacts with the gene’s promoter more strongly if it encodes the risk alleles. Based on these findings, disease susceptibility SNPs at *5q15* likely do not confer disease susceptibility by alternative splicing, but by changing enhancer-promoter interactions of *ERAP2*.

The SNP rs2248374 is located at the 5’ end of the intron downstream of exon 10 of *ERAP2* within a donor splice region and strongly correlates with alternative splicing of precursor RNA (23, 26). While the A allele of rs2248374 results in constitutive splicing, the G allele is predicted to impair recognition of the motif by the spliceosome (**Figure 1A**), which is conceptually supported by reporter assays outside the context of the *ERAP2* gene (26). Through reciprocal SNP editing in genomic DNA, we here demonstrated that the genotype of rs2248374 determines the production of full-length ERAP2 transcripts and protein.

Exon 10 is extended due to the loss of the splice donor site controlled by rs2248374 and consequently includes premature termination codons (PTCs) embedded in intron 10-11. (23, 26). Transcripts that contain a PTC can in principle produce truncated proteins, but if translation terminates more than 50-55 nucleotides upstream (“50-55-nucleotide rule”) an exon-exon junction (51), they are generally degraded through a process called *nonsense-mediated mRNA decay* (NMD). Our data show that ERAP2 dramatically alter protein abundance proportionate to transcript levels, which is consistent with the notion that transcripts encoding the G allele of rs2248374 are subjected to NMD during steady-state (20, 23). The loss of ERAP2 is relatively unusual, given that changes in ERAP2 isoform usage manifest so dramatically at the proteome level (20, 52). However, that ERAP2 transcripts can escape NMD under inflammatory conditions, such that haplotypes that harbour the G allele of rs2248374 have been shown to produce truncated ERAP2 protein isoforms (29, 53), not to be confused with “short” ERAP2 protein isoforms that are presumably generated by post-translational autocatalysis unrelated to rs2248374 (54).

Most protein-coding genes express one dominant isoform (55), but since both alleles of rs2248374 are maintained at near equal frequencies (allele frequency ∼50%) in the human population, this leads to high interindividual variability in ERAP2 isoform profile (23). ERAP2 may enhance immune fitness through balanced selection, especially since recent evidence indicates that the presumed “null allele” (i.e., the G allele of rs2248374) encodes distinct protein isoforms in response to infection (29, 56). A recent and unusual natural selection pattern during the *Black Death* for the haplotypes tagged by rs2248374 (25) supports this.

Nowadays, these haplotypes also provide differential protection against respiratory infections (24), but they also modify the risk of modern autoimmune diseases like CD, BCR, and JIA. The SNP rs2248374 was long assumed to be primary responsible for other disease-associated SNPs near *ERAP2*. Using conditional association analysis and mechanistic data, we challenged this assumption by showing that autoimmune disease-risk SNPs identified by GWAS influence ERAP2 expression independently of rs2248374.

These findings are significant for two main reasons: First, these results demonstrate that chromosome structure plays important roles in the transcriptional control of *ERAP2* and thus that its expression is regulated by mechanisms beyond alternative splicing. We focused on a small *cis*-regulatory sequence downstream of *ERAP2* as a proof of principle. Here, we showed that disease-risk SNPs alter physical interactions with the promoter in immortalised lymphoblast cell lines from autoimmune patients and that substitution of the allele of one common SNP (rs2548224) significantly affected the expression levels of *ERAP2*.

Another significant reason is that these findings have implications for our understanding of diseases in which ERAP2 is implicated. We recognize that the considerable LD between SNPs near *ERAP2* indicates that the effects of rs2248374 on splicing, as well as other mechanisms for regulation (i.e., chromosomal spatial organisation), should often occur together. Because of their implications for the etiology of human diseases, it is still important to differentiate them functionally. In light of the fact that disease-associated SNPs affect ERAP2 expression independently of rs2248374, ERAP2 may be implicated in autoimmunity not because it is expressed in susceptible individuals but because it is expressed at higher levels (20, 31). It corresponds with the notion that pro-inflammatory cytokines, such as interferons, upregulate ERAP2 significantly, while regulatory cytokines, like TGF-β , downregulate it, or that ERAP2 is increased in lesions of autoimmune patients (56–58). Overexpression of ERAP2 may be exploited therapeutically by lowering its concentration in conjunction with local pharmacological inhibition of the enzymatic activity (59).

We do like to stress that results from conditional eQTL and pQTL analysis in this study, supported by data from chromosome conformation capture coupled with sequencing analysis (40), as well as MPRA data (32) suggest that more SNPs may act in concert to regulate *ERAP2* expression, illustrating how intricately ERAP2 is regulated. Additional experimental work is needed to interrogate the extended ERAP2 haplotype and follow up on some of the derived associations. Mapping all the putative functional implications of these SNPs by CRISPR-knockin experiments in genomic DNA is inefficient and labour-intensive, which makes their application in primary tissue challenging. MPRA provides a high-throughput solution to interrogating SNP effects, but lacks genomic context, and can only infer local allelic-dependent effects (i.e., no long-range interactions). Due to their dependency on PAM sequences for targeting regions of interest, CRISPR/Cas9-based enhancer-targeting systems (60) may not be able to dissect functional effects at a single nucleotide (i.e., SNP) resolution. It is possible to discern allelic-dependent effects in the canonical genomic context using allele-specific 4C sequencing, but in case of high LD and closely clustered SNPs (e.g., the ∼900 bp region identified in this study) functional or non-functional SNPs cannot be distinguished within the sequence window of interest. Regardless, by integrating information from all of these available technologies, we were able to shortlist an interval suitable for interrogation by CRISPR-based knock-in techniques. A major drawback of this multi-step approach is that our study is therefore limited by sample size, and ideally we should have successfully targeted the regulatory region in a larger number of cell lines. Also, while ERAP2 also shows tissue-shared genetic regulation, there may be important cell-type specific regulatory mechanisms enforced by disease risk allele, that require study of this mechanism in lesional tissues and under inflammatory conditions. Regardless, single-cell analysis shows that the many *ERAP2* eQTLs are shared between immune cells (47, 61). This indicates that the mechanism by which the SNPs in the *cis*-regulatory region increase ERAP2 promoter interaction may be ubiquitous.

An enhancer-promoter loop increases transcriptional output through complex organisation of chromatin, structural mediators, and transcription factors (62, 63). Although we narrowed down the *cis*-regulatory region to ∼900bp, the identity of the structural or transcriptional regulators that juxtapose this region with the *ERAP2* promoter remains elusive. Loop-forming transcription factors such as CTCF and protein analogous (e.g., YY1, the Mediator complex) have been shown to contribute o enhancer-promoter interactions (64–68). Given that the here identified *cis*-regulatory region is located within the *LNPEP* promoter, it is challenging to identify the factors responsible for ERAP2 expression, since promoters are highly enriched for a large variety of transcription factor footprints (i.e., high ChIP-seq signals). Further studies are required to dissect how these *ERAP2* eQTLs modify enhancer activity and transcription, and how these mechanisms are distinguished from canonical promoter activity for *LNPEP* gene.

In conclusion, these results show that clustered genetic association signals associated with diverse autoimmune conditions and lethal infections act in concert to control expression of *ERAP2* and demonstrate that disease-risk variants can convert a gene promoter region into a potent enhancer of a distal gene.

## Materials & methods

### Cell culture

The THP1 cell line (ATCC, TIB-202™ ECACC Cat# 88081201, RRID:CVCL_0006, monocyte isolated from peripheral blood from an acute monocytic leukaemia patient) and Jurkat cell line (ATCC, Clone E6-1,TIB-152™; established from the peripheral blood of a 14-year-old, male, acute T-cell leukaemia patient) were purchased from ATCC. Cell lines were cultured in Roswell Park Memorial Institute 1640 medium (RPMI 1640, Thermo Fisher Scientific) supplemented with 10% heat-inactivated foetal bovine serum (FBS, Biowest Riverside) and 1% penicillin/streptomycin (Thermo Fisher Scientific). The authenticity of each cell line was monitored by genome-wide SNP-array analysis (for technical details see *High-density SNP-array analysis* below). On the SNP array, 41 SNPs from a “97-SNP fingerprint” for cancer cell line authentication (COSMIC Cell Line repository, available via https://cancer.sanger.ac.uk) were present. Jurkat and THP-1 cell lines used in this study had genotypes identical to those in the COSMIC Cell Line repository (**Supplementary Table 1**).

### CRISPR-Cas9-mediated allelic substitution

We used the Alt-R gRNA system (Alt-R^®^ CRISPR-Cas9 tracrRNA ligated to custom Alt-R® CRISPR-Cas9 crRNA) together with recombinant Alt-R® S.p. Cas9 Nuclease V3 (Integrated DNA Technologies) and custom Ultramer® DNA Oligo (Integrated DNA Technologies) as donor templates to modify rs2248374 through homology directed repair (**Table 1**). The guide RNA with the recombinant Cas9 nuclease (RNP complex) was assembled by incubating the Alt-R^®^ tracrRNA with the custom crRNA (**Table 1**) at 95°C for 5 min (1:1 ratio), followed by cooling down at room temperature. The RNP complex was mixed with the Alt-R^®^ S.p. Cas9 Nuclease and Buffer R (Neon system), followed by 10 min incubation. The custom DNA template was added to the mixture after RNP assembly. THP-1 cells or Jurkat cells were electroporated with the Neon Transfection System (Thermo Fisher Scientific) (for Jurkat cells: protocol A in **Table 3**). After electroporation the cells were incubated overnight with antibiotic-free culture medium with 20 µM Alt-R® HDR Enhancer (Integrated DNA Technologies). The next day, cells were diluted in 10x dilution steps until a final concentration of 30 cells/mL was reached (total volume 10 mL). Cells were seeded in multiple flat-bottom 96-well plates at 20 µL cell suspension per well (<1 cell/well) and transferred to 24 well plates once grown confluently. Confluent cultured clones were lysed (∼80% of total volume) in RLT buffer (Qiagen, Cat# 1030963) and DNA was isolated using Qiagen AllPrep DNA/RNA/miRNA Universal Kit (Cat# 80224). The flanking sequence of rs2248374 was amplified by PCR (Forward primer: AGGGAAAGAGAAGAATTGGA; Reverse primer: TCTCTTTCCTGTAGTGATTC) and PCR products incubated with the TaqI-v2 (R0149S, New England Biolabs) restriction enzyme (15 min at 65 °C). Next, the PCR products were loaded on a 1% agarose gel (agarose, Acros Organics; 10x TAE UltraPure^TM^, Invitrogen) to assess restriction by TaqI. Samples with a cleaved PCR product were prepared for sanger sequencing to validate in-frame integration of donor DNA and correct modification of rs2248374. CRISPR-Cas9-mediated haplotype substitution (i.e., disease-risk alleles to protective alleles) was achieved by using a large single-strand DNA template (**Table 1b**, 1500 bp Megamer™, Integrated DNA Technologies) with the alternative allele for SNPs (from ‘5 to 3’) rs2548224, rs3842058, rs2548225, rs2617435, rs1046395, rs1046396, and rs2762 and two guide RNAs (assembled separately) targeting upstream of rs2548224 and downstream of rs2762 respectively (**Table 2**) to introduce two double strand breaks. The two RNP complexes were transfected together with the Megamer™ in Jurkat cells (**Table 3**, Jurkat protocol B). Jurkat cells were incubated overnight with antibiotic-free culture medium with 20 µM Alt-R® HDR Enhancer (Integrated DNA Technologies), followed by single cell seeding. PCR (Forward primer: GTGGGCAGTGGGAAAGTTGG; Reverse primer: TGTCTCCAGCATCAACTCTGA) and restriction analysis by TaqI (TaqI-v2, cleaves PCR products containing the G allele of rs1046396 tagging the protective allele) was used to identify clones without cleaved PCR bands (i.e., loss of risk haplotype), and allelic substitutions were validated by sanger sequencing.

**Table 1a:**
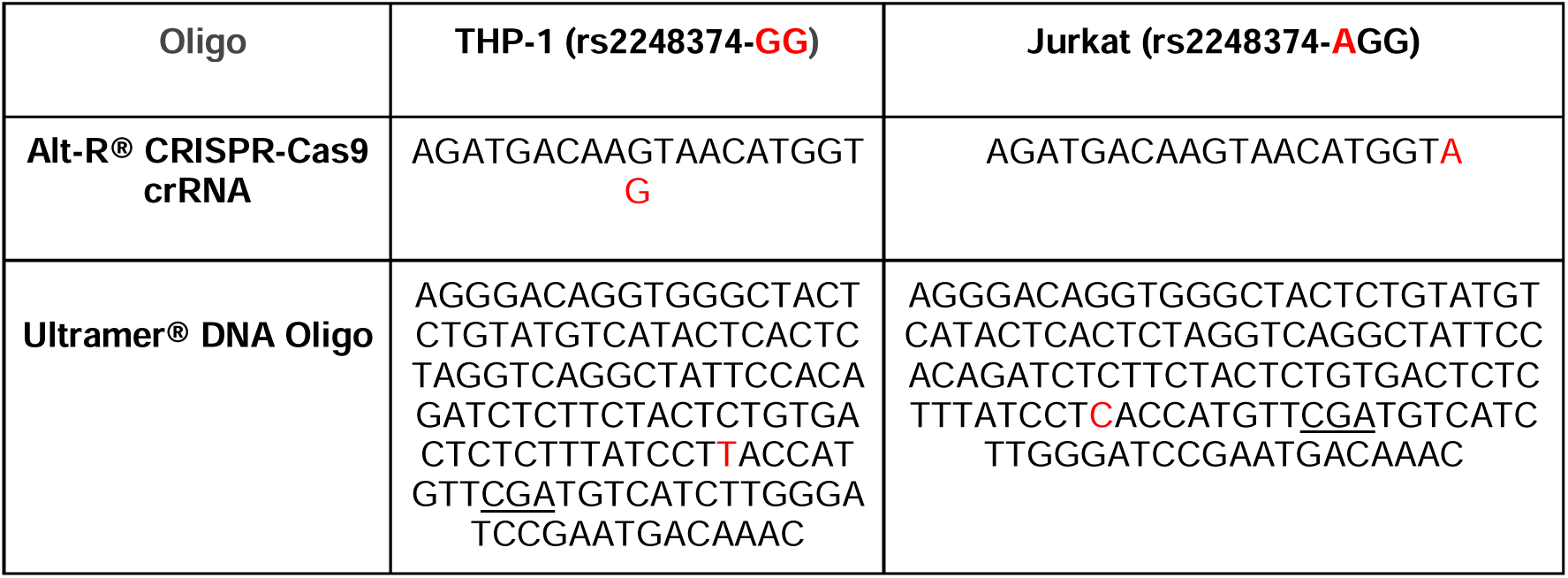
DNA oligonucleotides used as guide RNA and DNA donor template for modification of rs2248374 in both THP-1 and Jurkat cell lines. Sequences (5’→3’) of the used Alt-R® CRISPR-Cas9 crRNAs and Ultramer® DNA Oligos. Rs2248374 is highlighted in red in both crRNAs and DNA templates; the introduced *TaqI* motif sequence in each donor template is underscored (CGA). The template design of the Ultramer DNA oligos follows the work by Richardson *et al*. (69), which proposed that a template should be 127 bp in length (36 bp 5’, 91 bp 3’ of cut site respectively).

**Table 1b:**
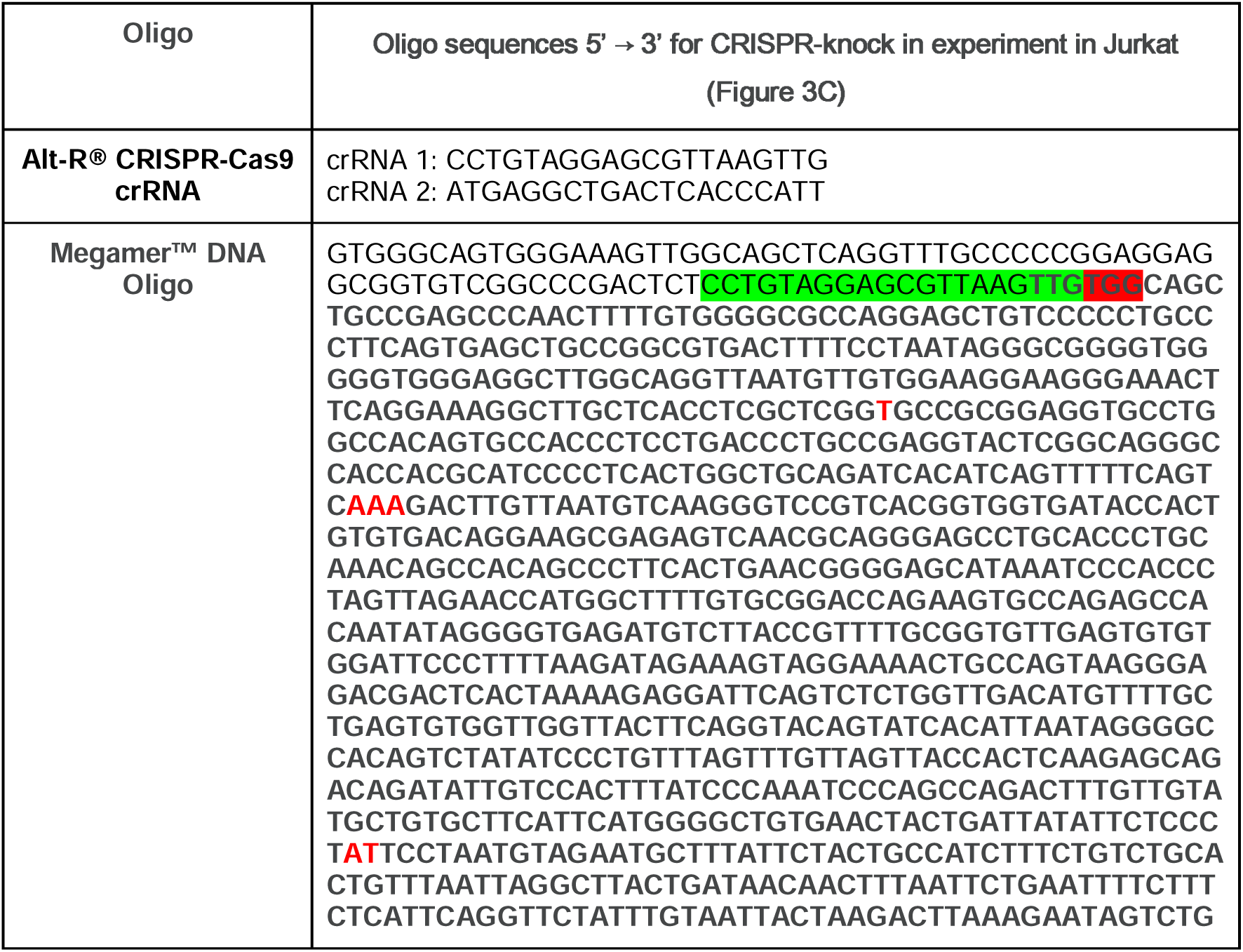
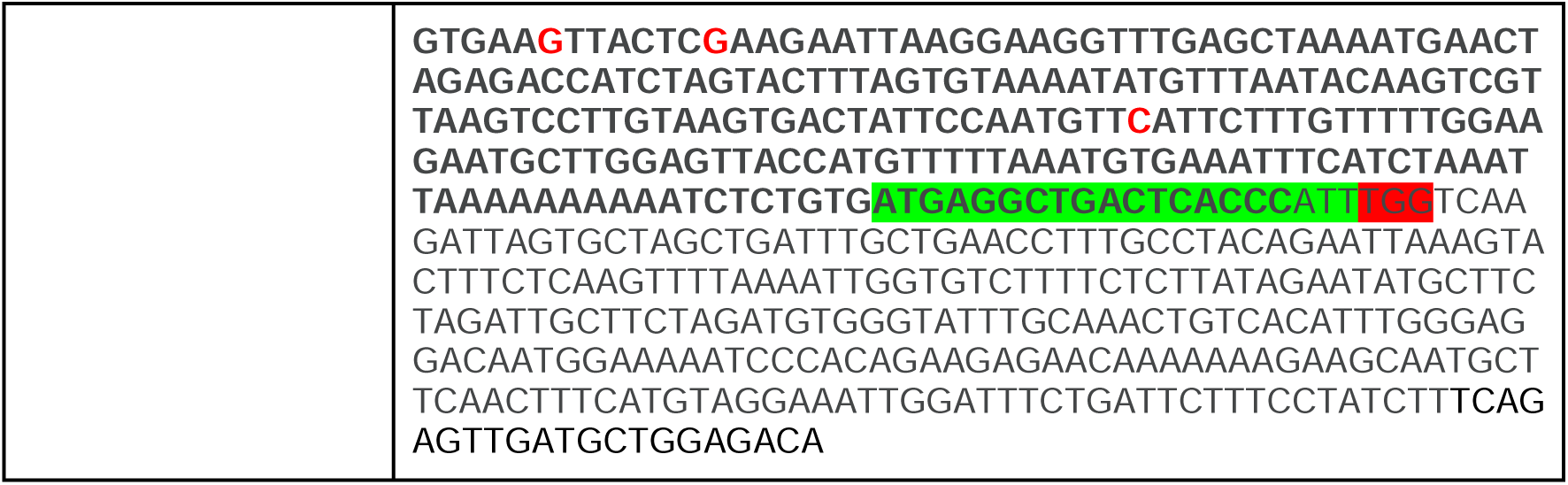
DNA oligonucleotides used as guide RNA and DNA donor template (1500bp) for modification of LNPEP promoter. Sequences (5’→3’) of the used Alt-R® CRISPR-Cas9 crRNAs and Megamer™ DNA Oligo. The protective alleles of the seven SNPs in the regulatory region are highlighted in red (from ‘5 to 3’) rs2548224-T, rs3842058-AAA, rs2548225-A, rs2617435-T, rs1046395-G, rs1046396-G, and rs2762-C respectively , in the Megamer™ sequence, as well as both gRNA binding locations (green = crRNA sequence; red = PAM sequence).

**Table 2.**
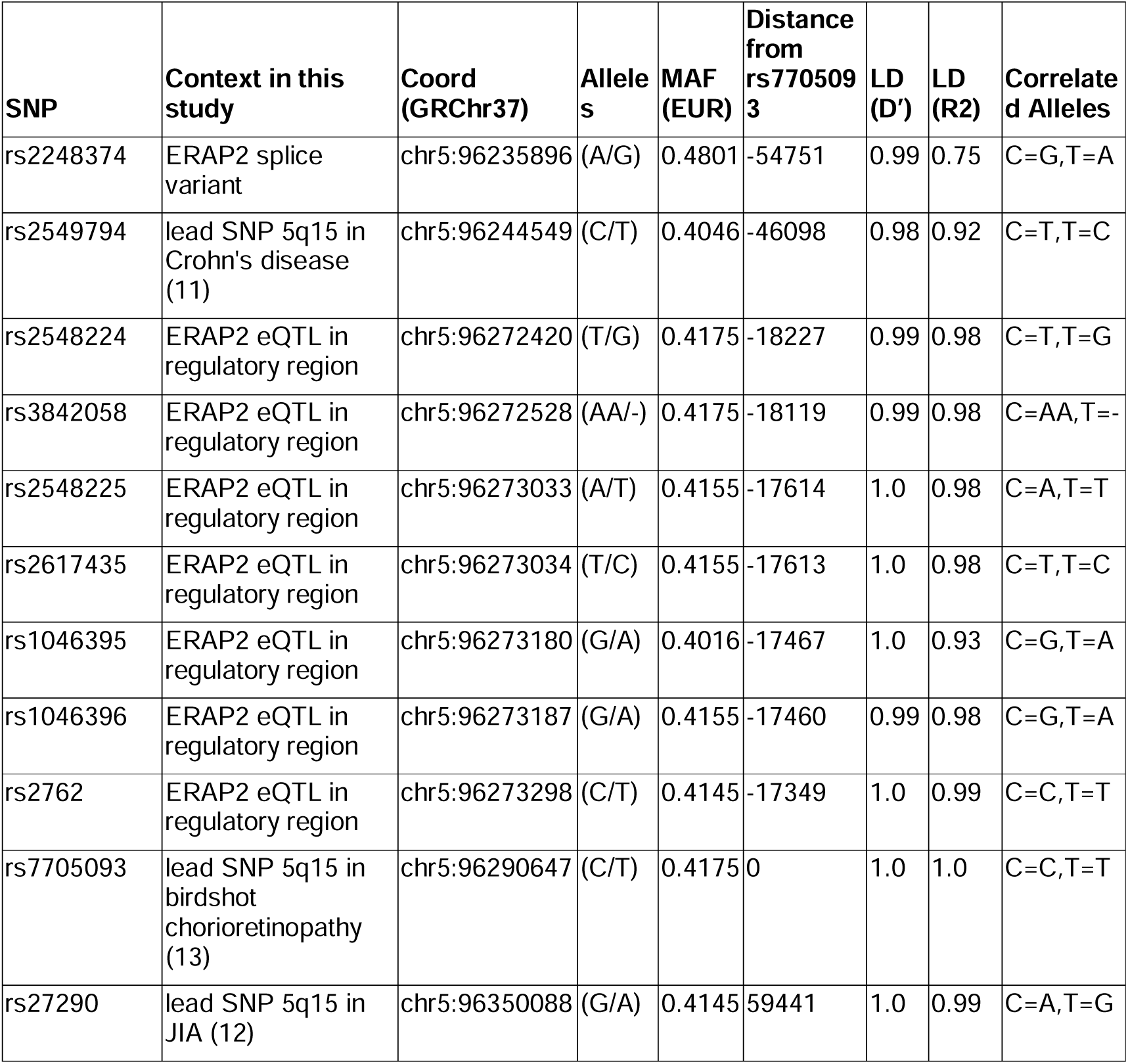
Details on the SNPs investigated in this study. The minor allele frequency (MAF) and linkage disequilibrium (LD) for each SNP is indicated for the European [EUR] superpopulation of the 1000 Genomes. Data was obtained from LDlink (70).

| SNP | Context in this study | Coord (GRChr37) | Alleles | MAF (EUR) | Distance from rs7705093 | LD (D') | LD (R2) | Correlated Alleles |
| --- | --- | --- | --- | --- | --- | --- | --- | --- |
| rs2248374 | ERAP2 splice variant | chr5:96235896 | (A/G) | 0.4801 | -54751 | 0.99 | 0.75 | C=G,T=A |
| rs2549794 | lead SNP 5q15 in Crohn's disease (11) | chr5:96244549 | (C/T) | 0.4046 | -46098 | 0.98 | 0.92 | C=T,T=C |
| rs2548224 | ERAP2 eQTL in regulatory region | chr5:96272420 | (T/G) | 0.4175 | -18227 | 0.99 | 0.98 | C=T,T=G |
| rs3842058 | ERAP2 eQTL in regulatory region | chr5:96272528 | (AA/-) | 0.4175 | -18119 | 0.99 | 0.98 | C=AA,T=- |
| rs2548225 | ERAP2 eQTL in regulatory region | chr5:96273033 | (A/T) | 0.4155 | -17614 | 1.0 | 0.98 | C=A,T=T |
| rs2617435 | ERAP2 eQTL in regulatory region | chr5:96273034 | (T/C) | 0.4155 | -17613 | 1.0 | 0.98 | C=T,T=C |
| rs1046395 | ERAP2 eQTL in regulatory region | chr5:96273180 | (G/A) | 0.4016 | -17467 | 1.0 | 0.93 | C=G,T=A |
| rs1046396 | ERAP2 eQTL in regulatory region | chr5:96273187 | (G/A) | 0.4155 | -17460 | 0.99 | 0.98 | C=G,T=A |
| rs2762 | ERAP2 eQTL in regulatory region | chr5:96273298 | (C/T) | 0.4145 | -17349 | 1.0 | 0.99 | C=C,T=T |
| rs7705093 | lead SNP 5q15 in birdshot chorioretinopathy (13) | chr5:96290647 | (C/T) | 0.4175 | 0 | 1.0 | 1.0 | C=C,T=T |
| rs27290 | lead SNP 5q15 in JIA (12) | chr5:96350088 | (G/A) | 0.4145 | 59441 | 1.0 | 0.99 | C=A,T=G |

**Table 3:** Settings used for transfection with the Neon Transfection System.

| Cell type | Cell # | Pulse (V) | Pulse width (ms) | Pulse # | Pipet tip type (μL) |
| --- | --- | --- | --- | --- | --- |
| THP-1 | $3.0 \times 10^5$ | 1600 | 10 | 3 | 10 |
| Jurkat (Protocol A) | $2 \times 10^5$ | 1100 | 30 | 2 | 10 |
| Jurkat (Protocol B) | $2 \times 10^5$ | 1325 | 10 | 3 | 10 |

### *CRISPR-Cas9* knock-out of *ERAP2* eQTLs

To delete a 116 kb region downstream of *ERAP2* in Jurkat cells, two RNP complexes were assembled in parallel using Alt-R gRNA system (Alt-R^®^ CRISPR-Cas9 tracrRNA ligated to custom Alt-R® CRISPR-Cas9 crRNA) together with recombinant Alt-R® S.p. Cas9 Nuclease V3 (Integrated DNA Technologies). RNPs with guide RNAs (guide RNA1; GGCATTCCTTAAGGGTATCA and guide RNA2; GCGTTGCTTCACATATAAGT, see for details **Supplemental Figure 1**) were transfected by electroporation into Jurkat cells (Jurkat protocol B in **Table 3**). Following transfection, Jurkat cells were diluted to 16.67 cells/mL and seeded in 96 well flat bottom plates (<1 cell/well at 50 µL cell suspension/well) for single cell cultures. Jurkat clones were screened by PCR (Fw: GTCCTTTCGCTGCTGATTTG; Rv: AGGTCATTCCACCACTTCATTGT), The PCR products were loaded on a 1% agarose gel (agarose, Acros Organics; 10x TAE UltraPure^TM^, Invitrogen) with the anticipated PCR product size only detectable after deletion of the entire DNA region, which was verified by sanger sequencing (**Supplemental Figure 1**).

### Western Blot analysis

Western blotting was used to determine the protein levels of ERAP2. NP40 lysis buffer was used to prepare total cell lysates (1% NP40, 135 mM NaCl, 5 mM EDTA, 20 mM Tris-HCl, pH = 7.4), complemented with cOmplete protease inhibitor cocktail (Roche). Protein lysates (20 μg/lane) were separated on a 4-20% Mini-PROTEAN TGX gel (Bio-Rad Laboratories) and transferred to a polyvinylidene difluoride membrane (Immobilon-P PVDF, Millipore). After blocking in 5% non-fat dry milk in TBST, membranes were probed overnight at 4°C with antibodies that recognize ERAP2 (AF3830, R&D Systems) or α-tubulin (T9026, Sigma-Aldrich). Following washing, membranes were incubated with either anti-goat or anti-mouse secondary antibodies conjugated to HRP (DAKO). Protein bands were detected with Amersham ECL™ Prime Western Blotting (RPN2236, GE Healthcare) on the ChemiDoc Gel Imaging System (Bio-Rad Laboratories).

### qPCR analysis

Gene expression was quantified by using qPCR. For *ERAP2* we used primers (**Table 4**) which bind to exon 10 (Fw: CATTCGGATCCCAAGATGAC) and exon 11 (Rv: GGAGTGAACACCCGTCTTGT) to determine functional *ERAP2* transcript based on rs2248374 variant. Primers targeting the control gene *RPL32* were used to calculate relative expression. (Fw: AGGGTTCGTAGAAGATTCAAGG; Rv: GGAAACATTGTGAGCGATCTC). The nucleotide sequence for the primers used for qPCR of *ERAP1* and *LNPEP* are shown in Table 4.

**Table 4:**
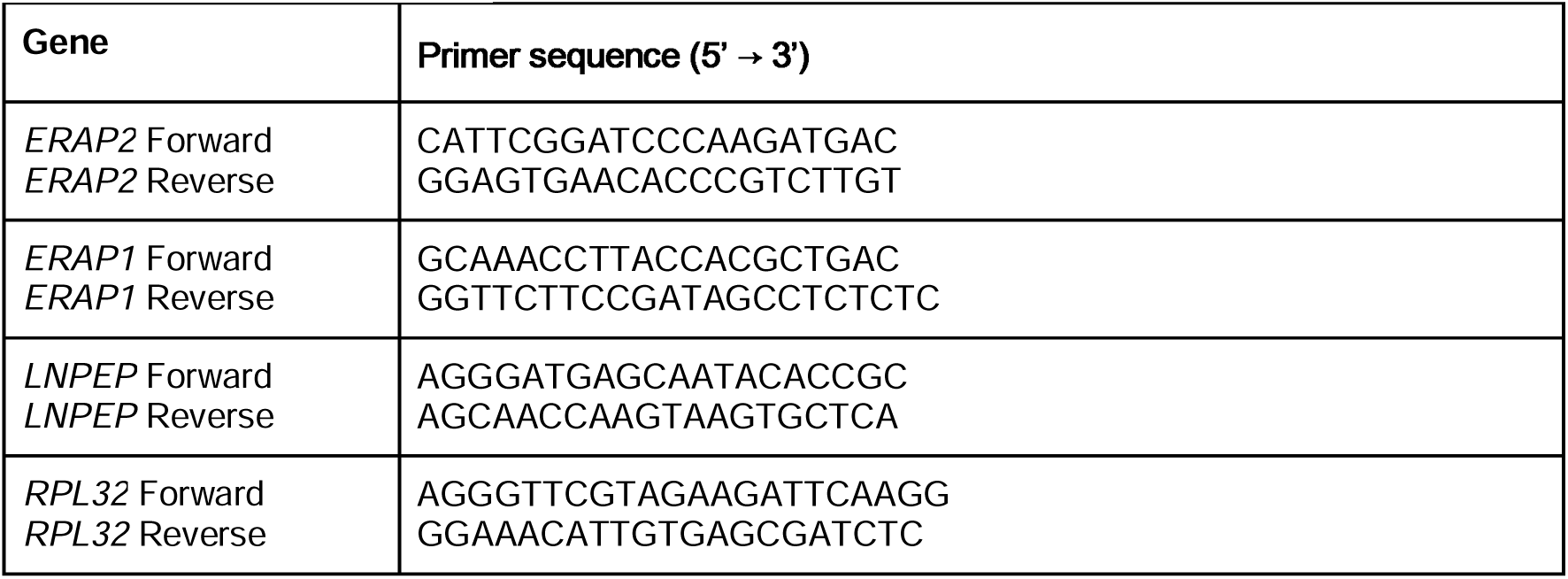
qPCR primers

### ERAP2 activity assay

To assess ERAP2 enzymatic function in THP-1 cells after introduction of the A allele of rs2248374, total cellular ERAP2 protein was enriched by immunoprecipitation and incubated with L-Arginine-7-amido-4-methylcoumarin hydrochloride (R-AMC, A2027, Sigma-Aldrich). Briefly, THP-1 cells were lysed (50 mM Tris, 150 mM NaCl, 1% Triton X-100, pH 7.5) and 500 µg cell lysate was incubated for 2h at 4°C with Protein-G-sepharose beads (BV-6511, BioVision) and anti-ERAP2 (AF3830, R&D Systems). Beads were washed in assay buffer (50 mM Tris, 1 mM DTT, pH 7.5) and resuspended in 100 µL assay buffer. Duplicate samples containing 40 µL beads solution were incubated with 40 µM of R-AMC (total volume 50 µL) for 1h at 37°C, while the fluorescent signal was measured (Ex: 370-10, Em: 440-20) with a CLARIOstar plate reader (BMG Labtech).

### Allele-specific 4C-seq

EBV-immortalised lymphoblastoid cell lines (LCL) were generated from peripheral blood mononuclear cells (PBMC) from 3 birdshot chorioretinopathy patients as previously described (33). The integrity of the genomes of the LCLs was assessed with whole genome SNP-array analysis, showing no abnormalities (**Supplemental Figure 2**). 4C template preparation was performed as described (34). Patient-derived cell lines were cultured in RPMI 1640 supplemented with 10% FBS 1% penicillin/streptomycin (Thermo Fisher Scientific), washed and cross-linked in 2% formaldehyde. DNA was digested in situ with MboI (NEB) and Csp6I (Thermo Scientific). Primers used for inverse PCR (5’- TACACGACGCTCTTCCGATCTTGTGGTTGGTTACTTCAGGT-3’, 5’- ACTGGAGTTCAGACGTGTGCTCTTCCGATCTGGACCCTTGACATTAACAAG-3’) allow reading of the ERAP2 eQTL rs3842058 (indel within the regulatory element) in the viewpoint fragment that is associated with the risk allele. Products were sequenced using Illumina sequencing (Illumina NextSeq 500). 4C-seq reads were demultiplexed by matching the 5’- ends of the R1 reads to the reading primer sequence (allowing 2 mismatches), and split into risk and non-risk reads based on the presence of the indel and its flanking sequence (GGACCCTTGACATTAACAAGTCTGAC for the risk allele and GGACCCTTGACATTAACAAGTCTTTG for the non-risk allele). Finally reads were mapped to the hg19 reference genome and processed using pipe4C (34) (github.com/deLaatLab/pipe4C) with the following parameters: normalization to 1 million reads in cis, window size 21, removal of top 2 read counts. Overlay plots were generated using R (https://www.R-project.org/).

### High-density SNP-array analysis

Genomic DNA of unedited and edited clones of THP-1 and Jurkat cell lines, and patient-derived LCLs was used for SNP-array copy number profiling and analysis of regions of homozygosity with the Infinium Human CytoSNP-850K v1.2 BeadChip (Illumina, San Diego, CA, USA). This array has ∼850,000 single nucleotide polymorphisms (SNPs) markers across the genome and can detect genomic insertions and deletions as well as stretches of homozygosity. Data analysis was conducted using NxClinical software v6.0 (Bionano genomics, San Diego, CA, USA). Human genome build Feb. 2009 GRCh37/hg19 was used. Results were classified with BENCH Lab CNV software (Agilent, Santa Clara, CA, USA). The genotype data were used for cell line verification and monitoring for potential genomic alterations in single cell cultures.

### ERAP2 eQTL colocalization and conditional analysis

Summary statistics of GWAS from CD (obtained via https://www.ibdgenetics.org/uploads/cd-meta.txt.gz) (11), JIA (12), and BCR (13) were used in visualisation of disease associated SNPs at *5q15*. ERAP2 eQTL data for ‘whole blood’ were downloaded from the GTEx portal and used for GWAS colocalization analysis with the *coloc* R package (v.5.1.0.1) (35). We assessed the likelihood of colocalization using the posterior probability (PP) for hypothesis 4 (that there is an association between both traits and they are driven by the same causal variant(s)). The likelihood of colocalization was deemed high for associations with PP4 > 0.8. The posterior probability of colocalization (H4) was calculated using the *coloc.abf()* function using default priors. For cross tissue ERAP2 eQTL conditional analyses, we used the “FastGxC” shared *ERAP2* eQTL data as reported by *Lu* et al., which is calculated using RNA-seq data from the Genotype-Tissue Expression (GTEx) Consortium v8 (36). To identify *ERAP2* eQTLs and pQTLs with association signals independent from rs2248374, conditional analysis was conducted using *conditional and joint multiple-SNP analysis* (GCTA-COJO) using the *ERAP2* eQTL data from FastGxC or ERAP2 pQTL data from targeted plasma proteomics from the INTERVAL study (37) using LD information from the EUR superpopulation as a reference panel. More details on colocalization and conditional association analysis are outlined in ***Appendix I***. The GWAS regional association data for CD, BCR, and JIA and ERAP2 eQTL and pQTL are shown in **Supplementary Table 2-7**.

### Splice prediction of rs2248374

We used the deep neural network *SpliceAI* (38) and *Pangolin* (39) to predict the effects of the A>G allele substitution in pre-mRNA transcript sequences. Masked scores were predicted via https://spliceailookup.broadinstitute.org/ ^38^ using "chr5 96900192 A G" as input for hg38, and "chr5 96235896 G A" for hg19.

### ERAP2 promotor-interacting SNP selection

The promoter-interacting eQTL summary statistics for *ERAP2* determined by H3K27ac HiChIP assays in primary immune cells were obtained from Chandra *et al.* (40). We selected *ERAP2* eQTLs located in active enhancer regions at *5q15* (i.e., H3K27ac peaks) that significantly interacted with the transcriptional start site of *ERAP2* for each immune cell type (*P*<0.05). We visualised *H3K27* acetylation data from primary tissue analysis from ENCODE using the *WASHU Epigenome Browser* (41) (v54.00) at http://epigenomegateway.wustl.edu/browser/ (direct link to used data at *5q15* via: https://tiny.one/2p92f7tb). Massive parallel sequence data results for *ERAP2* eQTLs were obtained from supplementary data from Abell *et al*. (32).

### Data availability

Additional data underlying figures is shown in **Supplemental Table 1-11.** The full reproducible code will be available via dataverseNL. https://dataverse.nl/

## Abbreviations

4C-seq: circular chromosome conformation capture
BCR: Birdshot chorioretinopathy
Cas9: CRISPR-associated 9 (nuclease)
CD: Crohn’s disease
CRISPR: Clustered Regularly Interspaced Short Palindromic Repeats
crRNA: CRISPR-RNA
CTCF: CCCTC-binding factor (transcription factor protein)
DNA: Deoxyribonucleic acid
EBV: Epstein-Barr virus
eQTL: expression quantitative trait loci
ERAP1: endoplasmic reticulum aminopeptidase 1
ERAP2: endoplasmic reticulum aminopeptidase 2
gRNA: guide-RNA
GWAS: genome-wide association study
H3K27ac: histone H3 lysine 27 acetylation
HDR: homology directed repair
HRP: horseradish peroxidase
JIA: juvenile idiopathic arthritis
LCL: lymphoblastoid cell
LD: linkage disequilibrium
LNPEP: Leucyl/cystinyl aminopeptidase
MHC-I: major histocompatibility complex class 1
MPRA: massively parallel reporter assay
mRNA: messenger ribonucleic acid
NHEJ: non-homologous end joining
NMD: nonsense-mediated decay
PAM: protospacer adjacent motif
PBMC: peripheral blood mononuclear cell
PCR: polymerase chain reaction
POL2RA: DNA-directed RNA polymerase II subunit RPB1
pQTL: protein quantitative trait loci
PTC: premature termination codon
qPCR: quantitative PCR
R-AMC: L-Arginine-7-amido-4-methylcoumarin hydrochloride
RNA: ribonucleic acid
RNA-seq: RNA sequencing
RNP: ribonucleoprotein
RPL32: ribosomal protein L32
SNP: single nucleotide polymorphism
TGF-β: transforming growth factor beta
tracrRNA: trans activating CRISPR-RNA
WT: wild-type
YY1: Yin Yang 1 (transcriptional repressor protein)

## Acknowledgements

We thank Dr. Dennis C. Ko, Dr. Darragh Duffy, and Dr. Jimmie Ye for helpful discussions.

## Supplemental figures

**Supplemental Figure 1:**
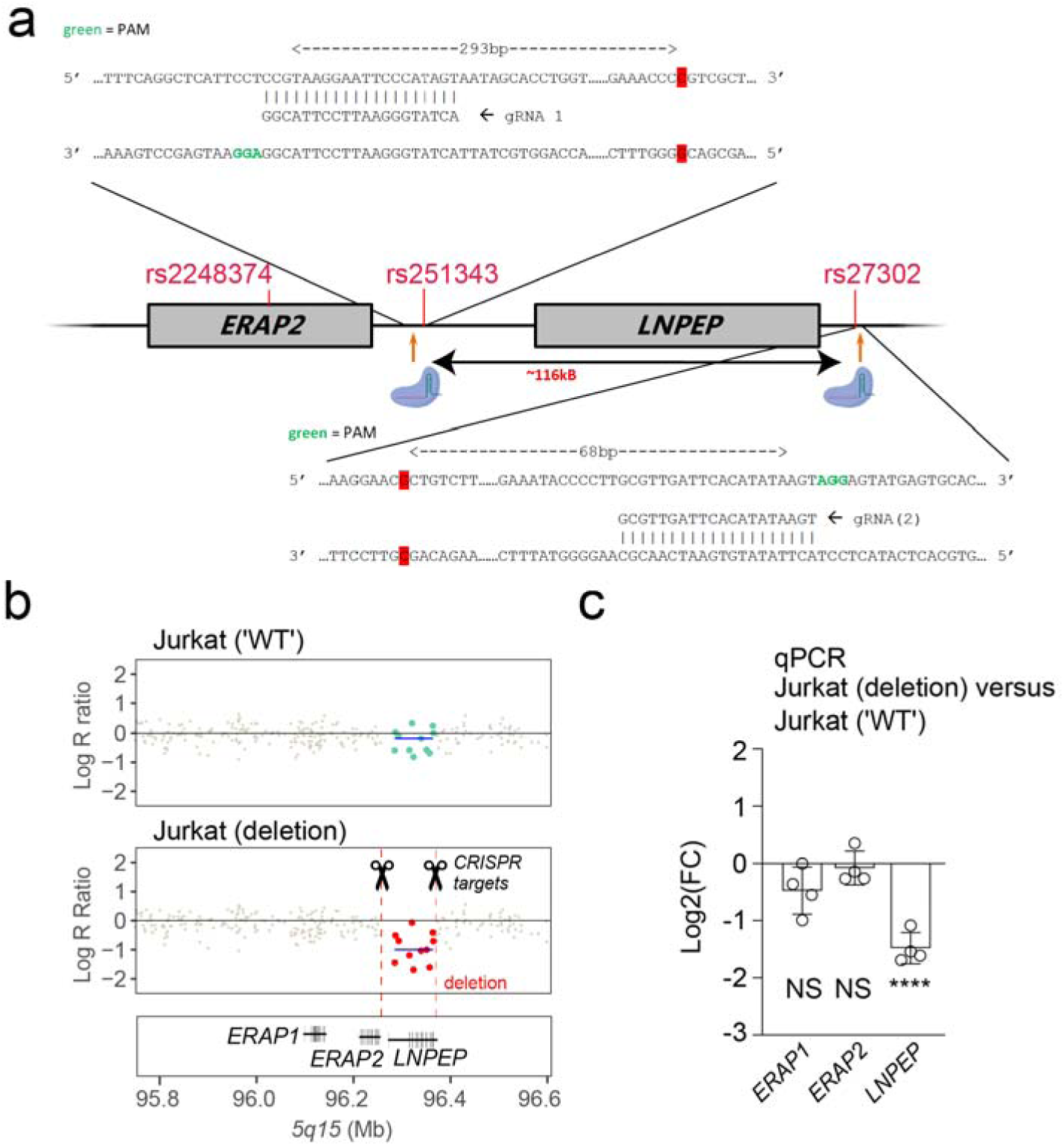
Deletion of downstream ERAP2 eQTLs using CRISPR Cas9. **A)** Overview of the guide RNA binding sites used for deletion of a 116 kb fragment that includes (from 5’ to 3’ direction) ERAP2 eQTLs from rs251343 until rs27302. Because Jurkat cells are heterozygous for rs10044354 located within the *LNPEP* gene (*P value, ERAP2 eQTL analysis* conditioned on rs2248374 = 3.2 x 10^-120^), we screened single cell cultures for specific deletion of rs10044354 by PCR. Application of this approach to Jurkat cells resulted in an clone with evidence for deletion at *5q15*, as confirmed by sanger sequencing and **B**) whole genome homozygosity SNP-array copy number profiling and analysis of regions of homozygosity using the Illumina CytoSNP-850K Beadchip. SNPs that reside in the region targeted by the CRISPR approach for deletion are denoted (green in unedited cells and red in the cells subjected to CRISPR-Cas9 with guide RNAs targeting this region as shown in *a*). Loss of SNP signal (in red) indicates partial deletion of ∼116 kB area. Genome wide results are shown in **Supplemental Figure 7**. **C**). Expression of *ERAP2, ERAP1* and *LNPEP* after CRISPR-Cas9 deletion, relative to unedited Jurkat cells (Log2(FC)). Data represents n = 4 replicates, Two-tailed unpaired t-test (NS = non-significant, **** *P*<0.0001.

**Supplemental Figure 2:**
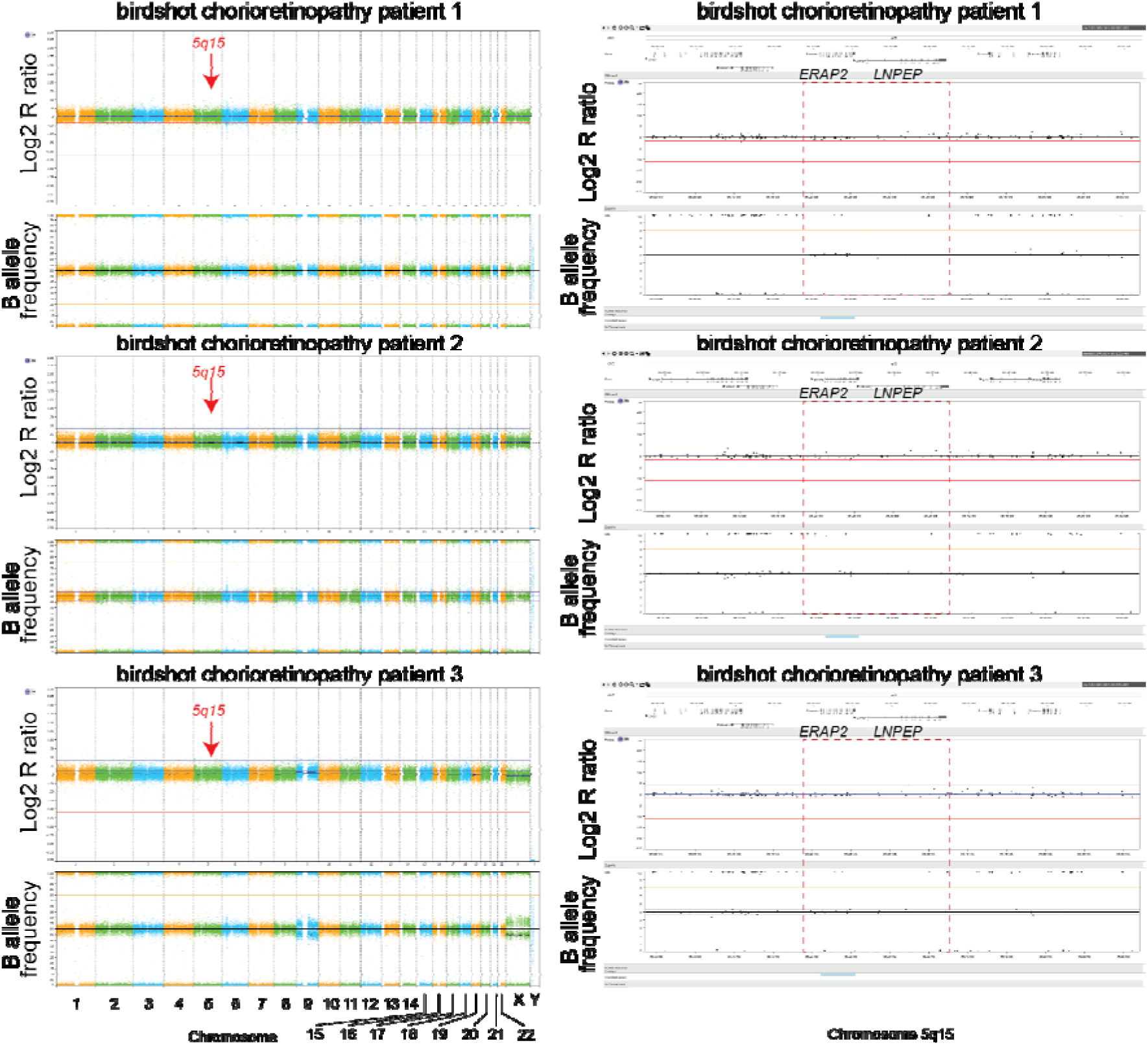
Whole genome analysis using SNP-arrays for birdshot chorioretinopathy patient-derived lymphoblastoid cell lines used for allele-specific 4C-seq. SNP-array based copy number profiling and analysis of regions of homozygosity using the Infinium Human CytoSNP-850K v1.2 BeadChip (Illumina, San Diego, CA, USA) showed no undesired genomic abnormalities, except for LCLs derived from patient 3, which exhibited a possible low level mosaicism for trisomy 9 (∼10%) and a comparable percentage of mosaicism for a monosomy of chromosome X (∼10%). Zoom plots on the right of chromosome 5q15 show normal copy number for SNPs at *5q15* and B allele frequency of ∼0.5 (i.e., heterozygous) for SNPs of interest in the extended ERAP2 haplotype. The panels show the array results for the whole genome. On the X-axis the chromosomes and chromosomal region are indicated. The upper Y-axis shows the Log2 R ratio and the lower Y-axis indicates the B allele frequency for each SNP.

**Supplemental Figure 3:**
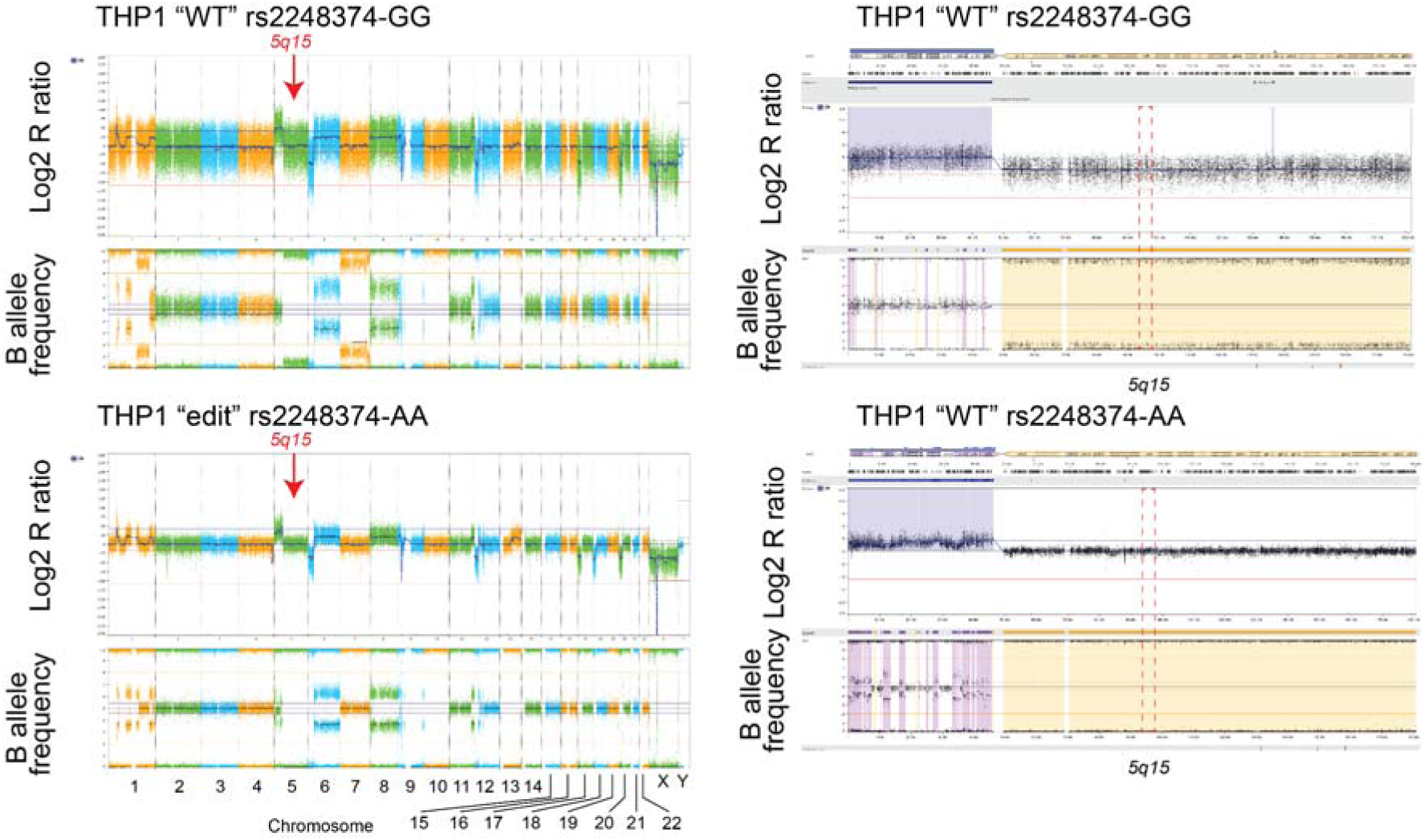
Whole genome copy number profiling and analysis of regions of homozygosity using SNP-arrays of the THP-1 cells used in this study. SNP-array based copy number profiling and analysis of regions of homozygosity using the Infinium Human CytoSNP-850K v1.2 BeadChip (Illumina, San Diego, CA, USA) showed multiple chromosomal abnormalities in the genome, including copy number neutral loss of heterozygosity (CN-LOH) for chromosome 5q, including the *5q15* genomic region. The panels show the array results for the whole genome on the right and for chromosome 5 on the left. On the X-axis the chromosomes and chromosomal region are indicated. The upper Y-axis shows the Log2 R ratio, and the lower Y-axis indicates the B allele frequency for each SNP.

**Supplemental Figure 4:**
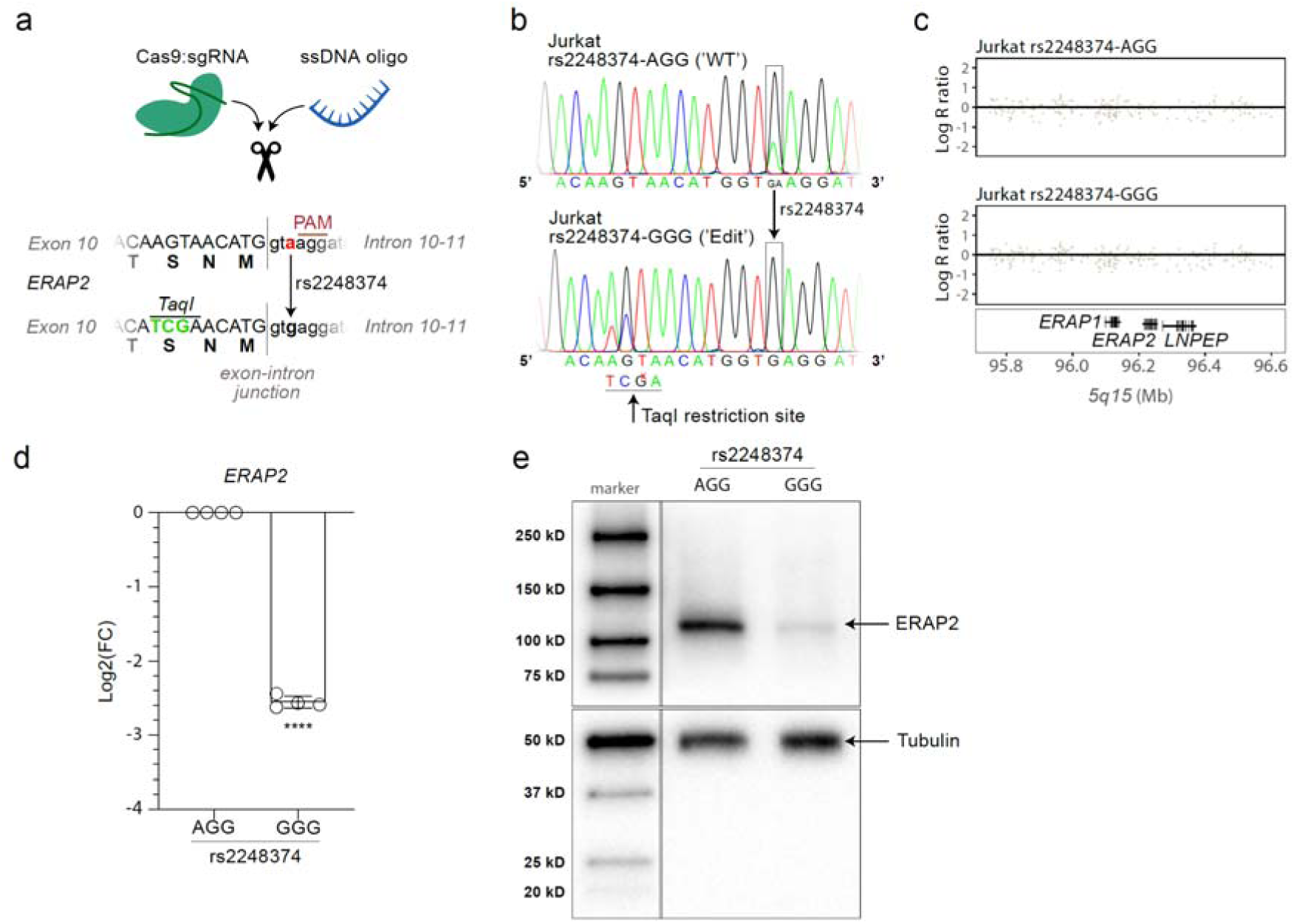
Allelic substitution of rs2248374 A>G allele abolishes ERAP2 expression. **A)** Overview of the homology-directed-repair (HDR) strategy to use CRISPR-Cas9 mediated SNP replacement of splice site variant rs2248374-AGG to GGG in Jurkat cells. An additional mutation was added to the ssDNA oligo template to introduce a TaqI restriction site, which was used for clone screening. **B**) Sanger sequence data showing Jurkat ‘WT’ with the heterozygous rs2248374-AGG variant and the Jurkat clone with the HDR-mediated SNP modification to rs2248374-GGG. **C**) Overview of SNP-array data, indicating no genomic abnormalities between unedited and edited clones at *5q15*. **D**) Decreased *ERAP2* expression was detected with qPCR as a result of the mutation of A to G allele of rs2248374. **E**) In addition, ERAP2 protein expression (determined by Western blot) was decreased in Jurkat cells that lost the A allele of rs2248374.

**Supplemental Figure 5.**
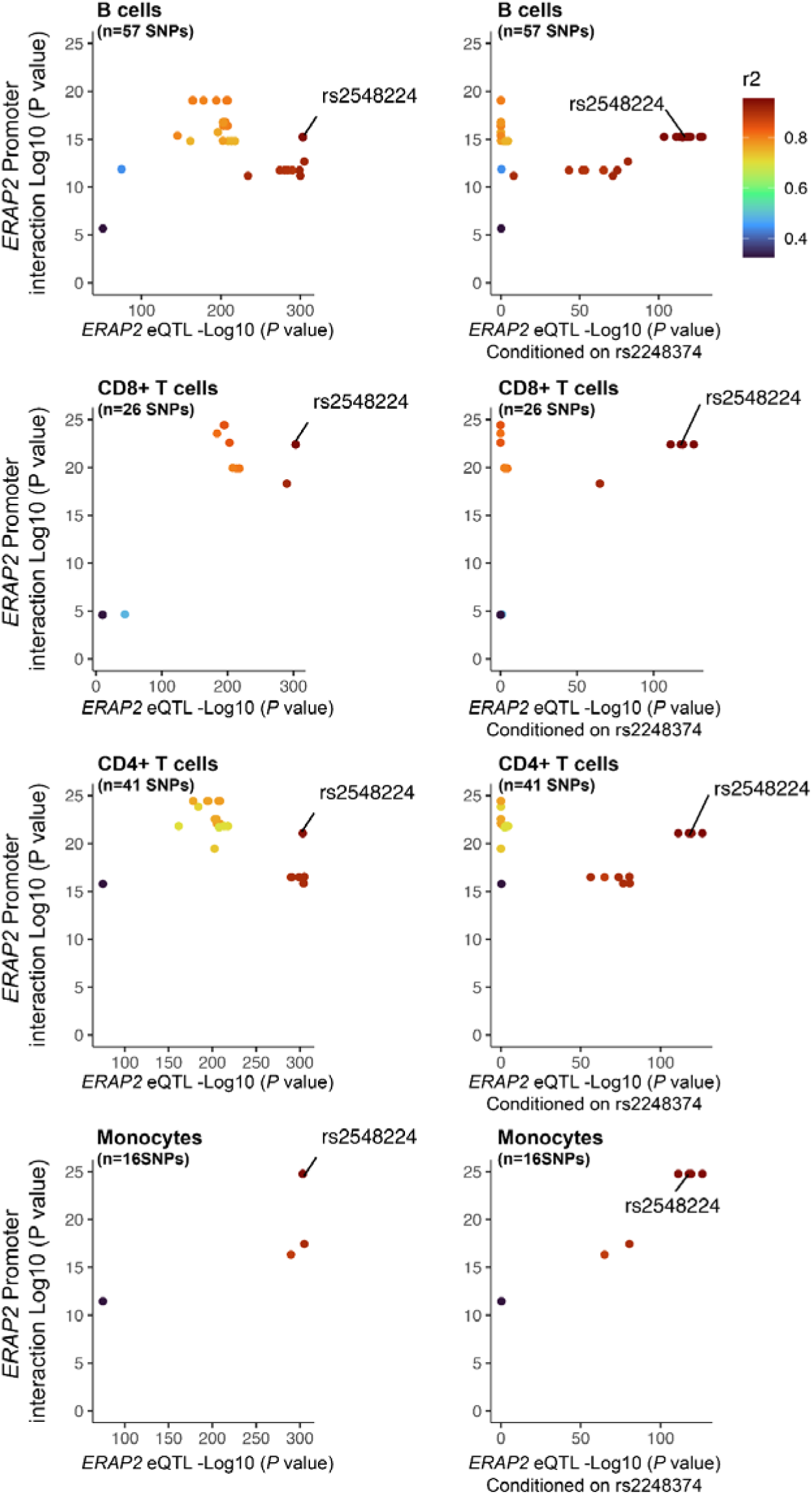
The SNPs rs2548224 is a strong *ERAP2* eQTLs independent of rs2248374 that reside in a DNA sequence that physically interacts with the *ERAP2* promoter. Comparison of the association analysis (*P* values) from chromosome conformation capture coupled with sequencing (Hi-C) data enriched by chromatin immunoprecipitation for the activating histone H3 lysine 27 acetylation (*H3K27ac*) in primary immune cells from Chandra *et al*. (40). SNPs with significant interaction (amount is indicated per plot) with *ERAP2* promoter in B cells, CD4+ T cells, CD8+ T cells and monocytes are shown. The colour intensity of each symbol reflects the extent of LD (r^2^ estimated using 1000 Genomes EUR samples) with rs2927608. The SNP rs2548224 that resides in a regulatory region is indicated.

**Supplemental Figure 6:**
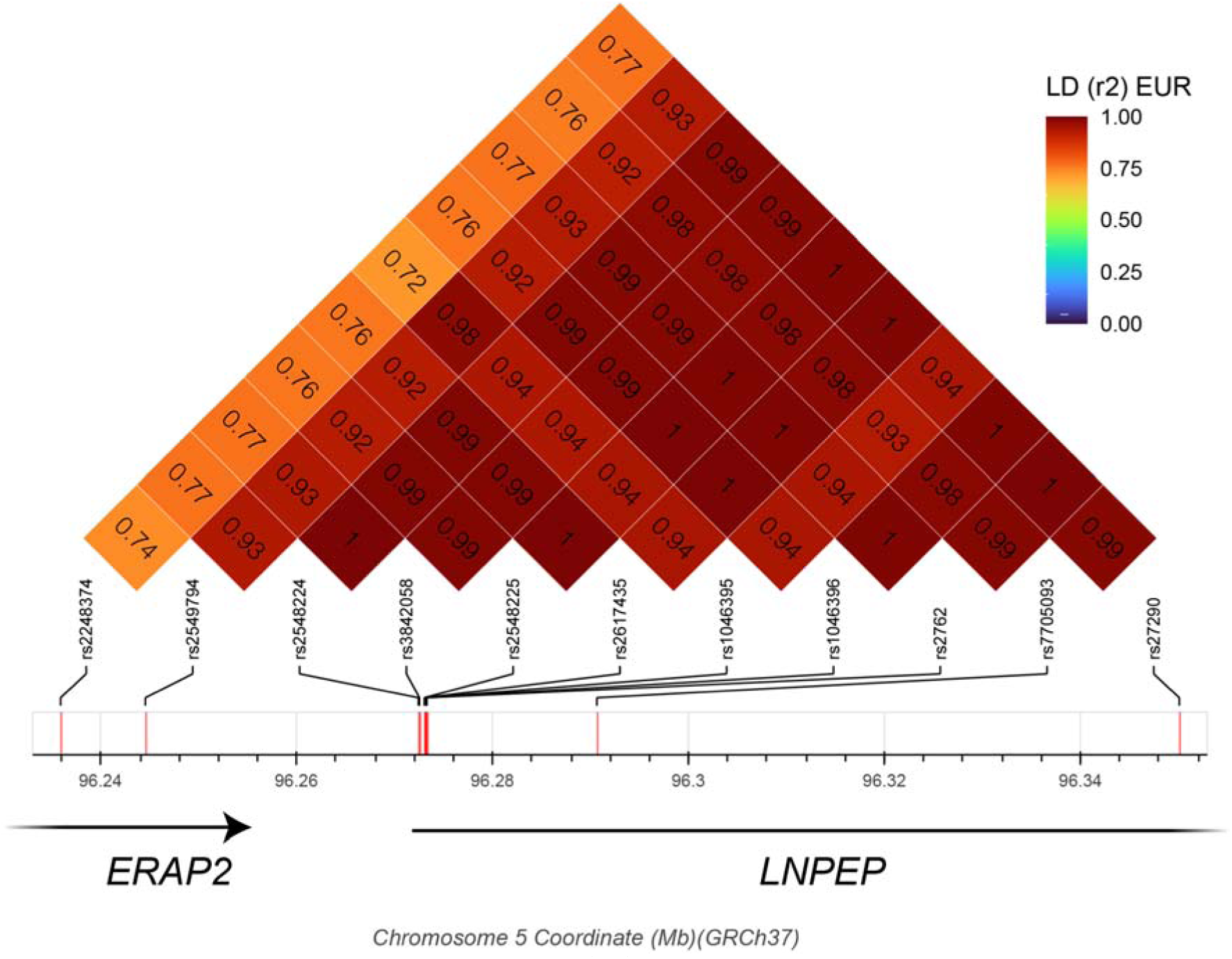
Linkage disequilibrium (LD) plot between disease-risk SNPs, rs2248374 and SNPs identified in Figure 3A.

**Supplemental Figure 7:**
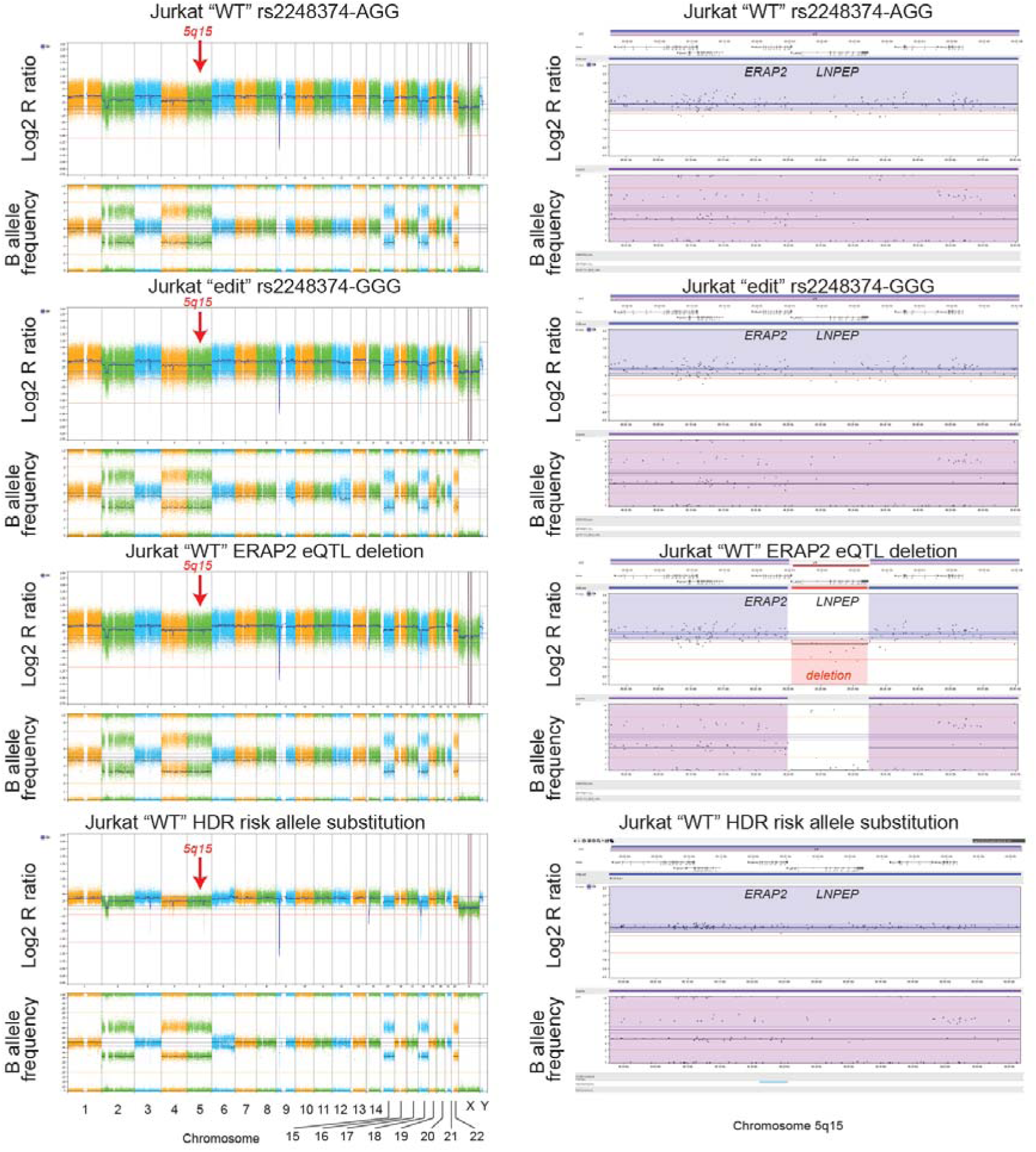
Whole genome copy number profiling and analysis of regions of homozygosity using SNP-arrays of the genome engineered Jurkat cell lines used in this study. SNP-array based copy number profiling and analysis of regions of homozygosity using the Infinium Human CytoSNP-850K v1.2 BeadChip (Illumina, San Diego, CA, USA) showed several genomic abnormalities in the Jurkat cell lines affecting multiple chromosomes, including a trisomy of chromosome 5. The copy number of the 5q15 genomic region was comparable between each condition, except for the intended deletion of a 116 kb fragment (in Jurkat “WT” *ERAP2* eQTL deletion). Zoom plots on the right for *5q15* with highlighted *ERAP2* and *LNPEP* genes. On the X-axis the chromosomes and chromosomal region are indicated. The upper Y-axis shows the Log2 R ratio and the lower Y-axis indicates the B allele frequency for each SNP.

**Supplemental Figure 8.**
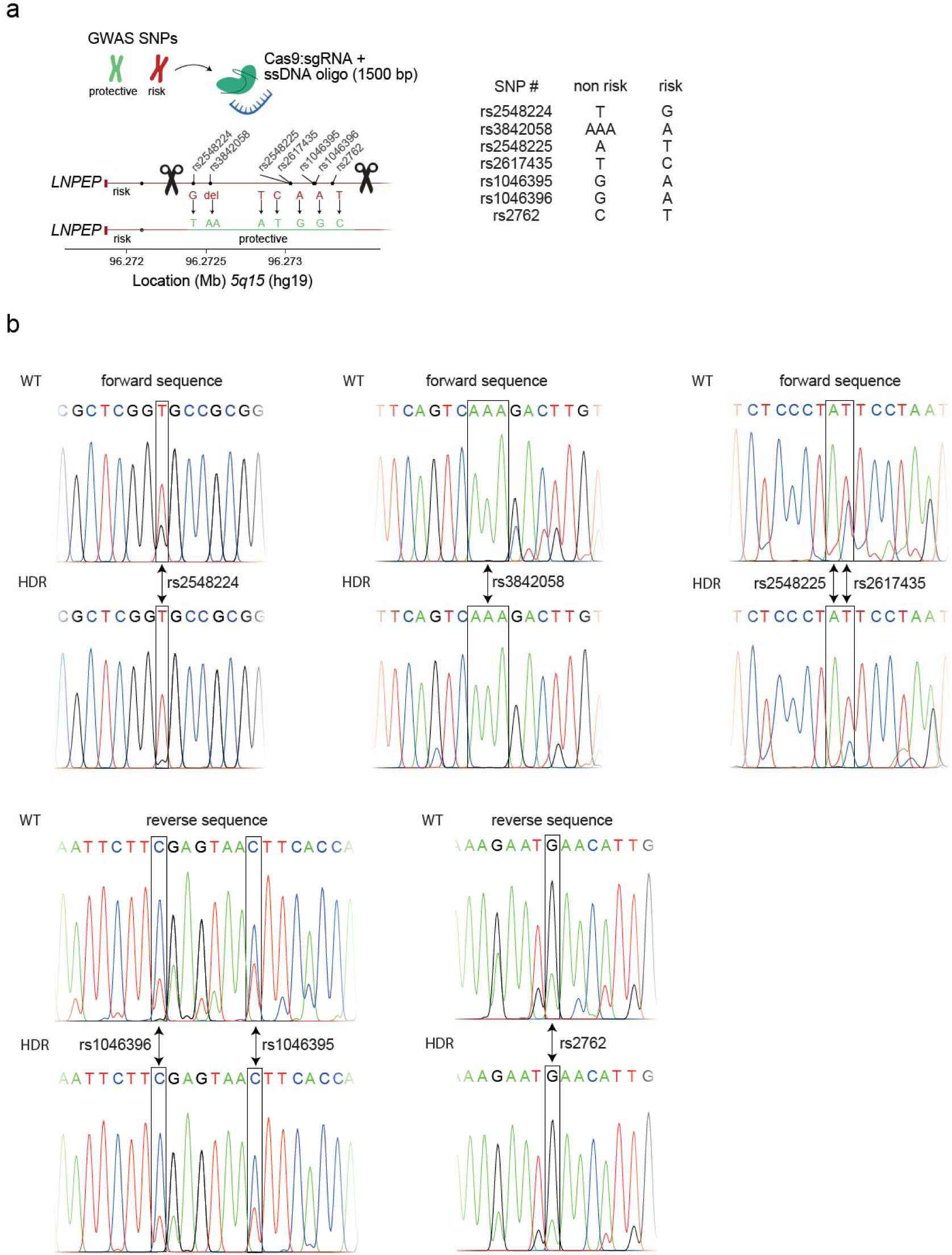
**A**) Overview of the homology-directed-repair (HDR) strategy to use CRISPR-Cas9 mediated SNP replacement in Jurkat cells to switch the alleles from disease risk SNPs (i.e., alleles associated with higher ERAP2 levels) to protective haplotype (i.e., alleles associated with lower ERAP2 expression). The region from 5’ to 3’ spans 879 bp. Note that Jurkat cells are triploid for chromosome 5 (e.g., for rs2548224 TTC > TTT would indicate a successful allelic substitution). **B**) Sanger sequencing results for the targeted Jurkat cells confirmed that the two cutting edges were accurately joined by HDR, and precisely switched alleles for rs2548224 leaving the haplotype otherwise intact.

## STAR Methods

### Key resource table

STAR Methods Key resource table
| REAGENT or RESOURCE | SOURCE | IDENTIFIER |
| --- | --- | --- |
| Antibodies |  |  |
| Goat polyclonal anti-LRAP/ERAP2 (1:2500) | R&D Systems | Cat# AF3830<br>RRID:AB_2099119 |
| Sheep polyclonal anti-LNPEP (1:2500) | R&D Systems | Cat# AF6386<br>RRID:AB_10717574 |
| Mouse monoclonal anti-α-Tubulin (clone DM1A) (1:5000) | Sigma-Aldrich | Cat# T9026<br>RRID:AB_477593 |
| Anti-CTCF |  |  |
| Chemicals, peptides, and recombinant proteins |  |  |
| Alt-R® HDR Enhancer | Integrated DNA Technologies | Cat# 1081072 |
| L-Arginine-7-amido-4-methylcoumarin hydrochloride | Sigma-Aldrich | Cat# A2027 |
| Amersham ECL™ Prime Western Blotting | GE Healthcare | Cat# RPN2236 |
| IGEPAL® CA-630 (NP40 substitute) | Sigma-Aldrich | Cat# I8896 |
| cOmplete Protease Inhibitor Cocktail | Roche | Cat# 11873580001 |
| Critical commercial assays |  |  |
| AllPrep® DNA/RNA/miRNA Universal Kit | Qiagen | Cat# 80224 |
| Infinium Human CytoSNP-850K v1.2 BeadChip | Illumina | Cat# 20025644 |
| Deposited data |  |  |
| qPCR data | This paper | Dataverse |
| R code for SNP-array plots | This paper | Dataverse |
| Experimental models: Cell lines |  |  |
| Human: THP-1 | ATCC | TIB-202™ |
| Human: Jurkat | ATCC | TIB-152™ |
| Human: Birdshot patient derived lymphoblastoid cells | This paper (see methods) | - |
| Oligonucleotides |  |  |
| qPCR primers, see Table 2 | Integrated DNA Technologies |  |
| Alt-R crRNAs, see Table 1A | Integrated DNA Technologies |  |
| Alt-R tracRNA | Integrated DNA Technologies |  |
| Ultramer® DNA Oligo: | Integrated DNA Technologies |  |
| Megamer™ ssDNA Fragment, see Table 1B | Integrated DNA Technologies |  |
| Software and algorithms |  |  |
| R Studio | R.4.2.2. |  |
| NxClinical software v6.0 | Bionano genomics |  |
| BENCH Lab CNV | Agilent |  |
| R package *sangerseqR* | Hill *et al.* (71) | https://www.bioconductor.org/packages/release/bioc/html/sangerseqR.html |
| Graphpad Prism | Graphpad Software Inc. | www.graphpad.com |

## Appendix I

### GWAS summary statistics and conditional analyses

Figure 2A-2D and 2F use the LD (r2) between SNPs at *5q15* rs2927608. This SNP is the top *ERAP2* eQTL (n=938) in the GTEx data for tissue "whole blood" (**Supplementary Table 2**). LD information for rs2927608 was estimated using the EUR superpopulation of 1000 Genomes in Plink 1.09 (http://pngu.mgh.harvard.edu/~purcell/plink/) (72). The genotype data for the EUR superpopulation (73) was extracted using the ensemble data slicer (74) with genotype file obtained via http://hgdownload.cse.ucsc.edu/gbdb/hg19/1000Genomes/phase3/ALL.chr5.phase3_shapeit2_mvncall_integrated_v5a.20130502.genotypes.vcf.gz and region lookup parameter "5:96000000-98000000". Sample ID’s from the EUR superpopulation of the 1000 Genomes were obtained by filtering using meta-data from the 1000 Genomes ftp sever url: http://ftp.1000genomes.ebi.ac.uk/vol1/ftp/release/20130502/integrated_call_samples_v3.20130502.ALL.panel and converted into a .vcf files using the *vcfR R* package and formatted in Plink 1.09 by the “--recode” option into a BED file. We calculated the allele frequency of each SNP using the --freq function in Plink 1.09 and LD with rs2927608 using the “--r2” option (11930 variants from position 96000022-97252405 with LD information available).

In the *Data availability* section we provide a link to the repository with the full GWAS summary statistics for BCR as reported by Kuiper *et al*. (13). The summary statistics from the BCR GWAS was filtered for variants at (Hg19) chromosome 5 (chr5) from position 96000000-97200000 (n=6307, **Supplementary Table 3**). For colocalization analysis, we cross referenced these BCR variants with GTEx v8 *ERAP2* eQTL from whole blood to obtain a final set of 679 SNPs. The CD GWAS summary statistics were downloaded from https://www.ibdgenetics.org/uploads/cd-meta.txt.gz and filtered for variants at (Hg18) chr5 from position 96081460-96461460 (n=197, **Supplementary Table 4**) of which 195 overlapped with GTEx *ERAP2* eQTL from whole blood for colocalization analyses. Because rs2248374 is not in the summary statistics of CD, we used the association statistics for variant rs2549782 (LD (r2) = 1.0 in EUR of 1000 Genomes). We obtained the GWAS summary statistics for JIA as reported by Hinks *et al.* (12). We filtered for variants at (Hg18) chr5 from position 95159342-98830661 (n=974, **Supplementary Table 5**) of which 495 variants were in common with the GTEx *ERAP2* eQTL data from whole blood and used in colocalization analysis. Note that we used the GWAS summary statistics reported by Hinks *et al*. that were calculated under a dominant genetic model.

To identify significant associations for SNPs with *ERAP2* mRNA and protein expression independent of the genotype of rs2248374, we performed approximate conditional analysis using GCTA v1.94.1 using the ‘--cojo-cond’ option (75). Our analysis uses the "FastGxC" tissue-shared *ERAP2* eQTL data for SNPs reported by Lu and associates, which is calculated using RNA-seq data from the GTEx Consortium GTEx v8 (46). For GCTA-COJO, we used linkage disequilibrium (LD) information from the EUR superpopulation (73) as a reference panel (see above, n=2903 SNPs were used after cross referencing with the 11930 variants from position 96000022-97252405 Hg19 from the EUR, **Supplementary Table 6**). The summary statistics for *ERAP2* pQTL data from blood plasma of the INTERVAL study as reported by Sun *et al*. (37) was downloaded from https://app.box.com/s/u3flbp13zjydegrxjb2uepagp1vb6bj2. We filtered for the variants near *ERAP2*, which resulted in 6001 pQTL used in approximate conditional analysis (**Supplementary Table 7**).

